# Dopamine signaling drives skin invasion by human-infective nematodes

**DOI:** 10.1101/2025.01.29.635547

**Authors:** Ruhi Patel, Aracely Garcia Romero, Astra S. Bryant, George W. Agak, Elissa A. Hallem

## Abstract

Skin-penetrating nematodes are one of the most prevalent causes of disease worldwide – nearly 15% of the global population is infected with at least one species of skin-penetrating nematode^1,2^. The World Health Organization has targeted these parasites for elimination by 2030^3^, but the lack of preventative measures is a major obstacle to this goal. The infective larvae of skin-penetrating nematodes enter hosts through skin^4^, and blocking skin penetration is an as-yet unexplored approach for preventing infection. However, in order to prevent worm ingress via the skin, an understanding of the behavioral and neural mechanisms that drive skin penetration is required. Here, we describe the skin-penetration behaviors of the human-infective threadworm *Strongyloides stercoralis*. Using fluorescently labeled worms to enable visualization on the skin coupled with time-lapse microscopy, we show that *S. stercoralis* engages in repeated cycles of pushing, puncturing, and crawling on the skin surface before penetrating the skin. Pharmacological inhibition of dopamine signaling inhibits these behaviors in *S. stercoralis* and the human hookworm *Ancylostoma ceylanicum,* suggesting a critical role for dopamine signaling in driving skin penetration across distantly related nematodes. CRISPR-mediated disruption of dopamine biosynthesis and chemogenetic silencing of dopaminergic neurons also inhibit skin penetration. Finally, inactivation of the TRPN channel TRP-4, which is expressed in the dopaminergic neurons, blocks skin penetration on both rat and human skin. Our results suggest that drugs targeting TRP-4 and other nematode-specific components of the dopaminergic pathway could be developed into topical prophylactics that block skin penetration, thereby preventing infections.

## INTRODUCTION

Skin-penetrating gastrointestinal parasitic nematodes, including the threadworm *Strongyloides stercoralis* and hookworms in the genera *Necator* and *Ancylostoma*, infect approximately one billion people worldwide and cause devastating disease and socioeconomic burden^1,2^. Infections by these parasites stunt development in children^5-7^, cause chronic disease in both children and adults^1,8,9^, and can be fatal for immunocompromised individuals^8,9^. Infections are most prevalent in communities that lack access to sanitation infrastructure and clean drinking water^8,9^, which perpetuates a cycle of socioeconomic disparity. Although drug treatments exist, these remedies do not prevent reinfection and may soon be rendered ineffectual by the evolution of anthelmintic-resistant nematode populations; indeed, anthelmintic resistance is a longstanding problem among nematodes that parasitize livestock^10^. Thus, there is an urgent need to expand the existing arsenal of medications to include preventative treatments.

Skin-penetrating nematodes infect hosts by penetrating through host skin^4^. As a crucial step of the infection process, skin penetration is a promising target for intervention – preventative treatments that block skin penetration would stop infections from establishing. Infective larvae are known to penetrate skin head-first^11^; however, beyond this, nothing is known about the behavioral strategies that are executed by skin-penetrating nematodes during skin invasion. The neural and molecular basis of skin-penetration behavior is also unknown. A mechanistic understanding of skin penetration could be harnessed to develop the first prophylactic anthelmintics^3^.

Here, we examine the skin-penetration behaviors of the human-infective nematode *S. stercoralis*^12^. We show that infective larvae engage in repeated cycles of pushing on the skin, puncturing the skin, and crawling on the skin before ultimately penetrating the skin. Initial penetration attempts are sometimes aborted; infective larvae then crawl to a new location and re-initiate penetration. Thus, infective larvae actively explore the skin surface before selecting a location to penetrate. Pharmacological inhibition of dopamine signaling impaired skin penetration in *S. stercoralis*, the distantly related human hookworm *Ancylostoma ceylanicum,* and the rat-infective nematode *Strongyloides ratti*, suggesting that dopamine signaling plays a conserved role in driving skin penetration across multiple species of skin-invading nematodes. Skin penetration was also inhibited by CRISPR/Cas9-mediated disruption of the dopamine biosynthesis gene *Ss-cat-2* and chemogenetic silencing of the dopaminergic neurons. Finally, we show that genetic inactivation of the *Ss-trp-4* gene, which encodes a nematode-specific TRPN channel, severely impairs skin penetration. Our results suggest that topical compounds that block *Ss*-TRP-4 or another nematode-specific component of the dopaminergic pathway could function as the first topical repellents for skin-penetrating nematodes.

## RESULTS

### *S. stercoralis* infective larvae engage in repeated behavioral cycles on skin

*S. stercoralis* penetrates human skin as developmentally arrested, infective third-stage larvae (iL3s)^13^. After host invasion, development resumes and the nematodes follow a complex life cycle that includes both intra-host and extra-host life stages^13,14^ (Fig. S1). To examine the behaviors of *S. stercoralis* iL3s on skin, we developed an *ex vivo* tracking assay that enabled us to observe and quantify skin penetration in real time (Fig. 1A). Briefly, we excised the skin from euthanized rats, removed the fur, sectioned the skin, and then suspended skin pieces over saline using plastic inserts. We then placed individual, fluorescent *S. stercoralis* iL3s on the surface of the skin and used a fluorescence dissection microscope and attached camera to acquire time-lapse images of worm behavior. The *S. stercoralis* iL3s were fluorescent across the entire body either because of stable expression of an *Ss-act-2p::strmScarlet-I* reporter cassette^15^ or because they were labeled with DiI (1,1’-dioctadecyl-3,3,3’,3’-tetramethylindocarbocyanine perchlorate), which stains the nematode cuticle^16^. This allowed visualization of the translucent worms on the skin surface.

**Figure 1.**
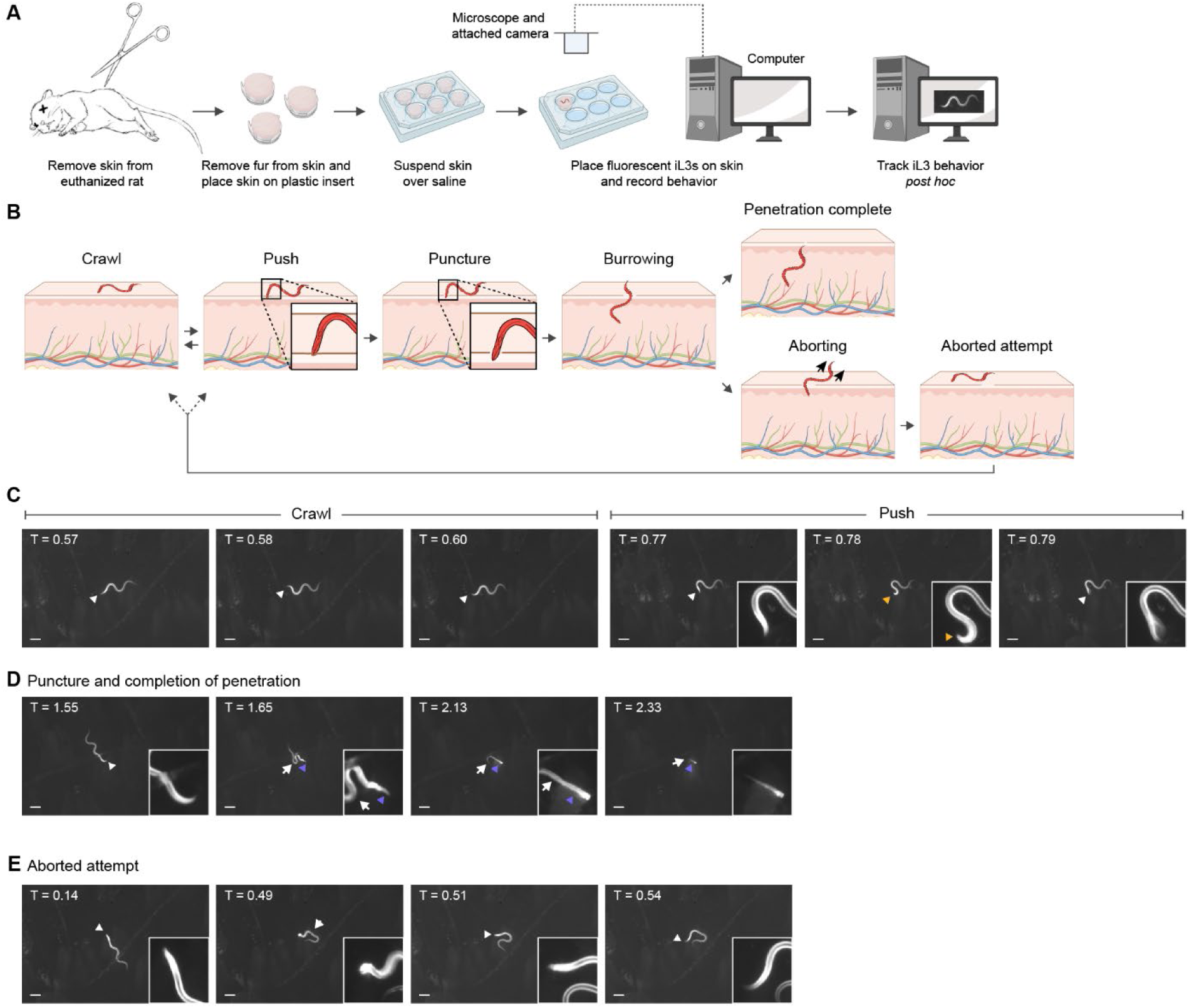
*S. stercoralis* iL3s engage in repeated behavioral motifs on skin. **A.** An *ex vivo* skin penetration assay was used to observe and quantify the behaviors of individual iL3s on the surface of rat skin. Skin was sourced from euthanized rats. The skin was excised from the rat and fur was manually removed from the skin surface. The skin was then sectioned into small pieces and the pieces were suspended over baths of BU saline^58^, using plastic cell-culture inserts, in individual wells of either a 6-well or 12-well plate. Next, fluorescent iL3s, which were fluorescent either because of dye-staining with DiI^16^ or because of expression of an *Ss-act-2p::strmScarlet-I* transgene^15^, were placed on the skin surface and time-lapse images were acquired for 5 min thereafter or until skin penetration was complete. Time-lapse images were recorded using a fluorescence dissection microscope and attached camera. The images were analyzed *post hoc* to quantify skin-penetration behaviors. **B.** Schematic depicts the behaviors executed by iL3s on skin. Crawling on the skin surface is characterized by sinusoidal locomotion. Pushes are characterized by pauses in locomotion, coupled with the worm moving its head back and forth, at an angle, against the skin. The head of the worm does not enter the skin during a push, and the epidermal surface remains intact (inset). Worms initiate a penetration attempt via a puncture, which is almost identical to a push, except that the head of the worm is detected inside the skin (inset). Thereafter, the worm continually burrows into the skin until penetration is complete, as characterized by full entry of the worm into the skin. In some cases, iL3s that are actively penetrating skin will abort the penetration attempt prior to completing penetration; these iL3s then crawl to a distinct spot, push, and subsequently penetrate the skin. The iL3s either abort the attempt by reversing completely out of the skin or by executing a turn within the skin and crawling out. **C-E.** Time-lapse images of an *S. stercoralis* iL3 engaging in skin penetration on rat skin. See also Movie S1. **C.** The iL3 crawling on the skin surface (left three panels) and pushing down on skin (right three panels). White arrowhead indicates the head of the worm. Yellow arrow indicates an instance of pushing, where the iL3 placed its nose almost perpendicular to the surface of the skin and pushed against it. The head of the worm appears blurry in the rightmost panel because it was actively pushing against the skin surface. **D.** The same iL3 puncturing and then penetrating skin. White arrowhead indicates the head of the worm, purple arrows indicate the part of the worm inside the skin, and white arrows indicate the part of the worm outside the skin. The first panel shows the iL3 outside the skin; the second panel shows that the iL3 has punctured the skin; the third panel shows that the worm has partially disappeared into the skin; and the fourth panel shows that the iL3 has almost completed skin penetration, with only the tip of the tail detectable outside. **E.** The same iL3 aborting an earlier attempt at penetration. The first panel shows the iL3 prior to puncturing the skin and the second panel shows that the iL3 has punctured the skin, with its head no longer detectable. The third and fourth panels show that the iL3 has aborted the penetration attempt and has returned to the skin surface; the entire iL3 is now visible again. White arrowhead indicates the head of the worm and white arrows indicate the part of the worm outside the skin. In panels C-E, the iL3 is shown in white and the skin is the darker surface beneath the worm. Timestamps listed at the top left corner of each panel are relative to the time of placement of the worm on skin in minutes. Scale bar = 100 µm and the inset shows a portion of the underlaid image magnified 4 times and adjusted to increase the contrast.

When *S. stercoralis* iL3s were placed on rat skin, they typically crawled a short distance (Fig. 1B-C, Movie S1) and then halted forward locomotion and pushed down perpendicularly on the skin surface with their heads (Fig. 1B-C, Movie S1). These pushes, where the head of the iL3 indented but did not pierce the skin (Fig. 1B-C), appeared to be a means of “sampling” the skin surface to identify soft spots or openings such as hair follicles that could be exploited for invasion. After pushing, iL3s either crawled to a distinct spot or initiated a skin-penetration attempt by puncturing the skin with their heads (Fig. 1B, D, Movie S1). Once a penetration attempt was initiated by a puncture, iL3s either continually burrowed into the skin until penetration was completed (*i.e.,* the full body of the iL3 was inside the skin, Fig. 1B, D), or they retreated to the skin surface and aborted the penetration attempt (Fig. 1B, E). Aborted penetration attempts may occur when iL3s encounter a structure within the skin that provides mechanical resistance to entry, such as the basement membrane, sebaceous glands, or fibroblasts^17^. Nearly all iL3s that aborted a penetration attempt subsequently re-initiated and completed penetration at a distinct site. Thus, skin penetration involves repeated sampling of the skin surface through cycles of pushing, puncturing, and crawling until the iL3 ultimately completes penetration. The finding that iL3s actively explore the skin surface to locate favorable entry points, rather than diving into the skin immediately upon contact, suggests that topical compounds that interfere with these behaviors could be developed into novel anti-nematode prophylactics.

### Penetration drive increases on host skin

To determine whether skin-penetration behaviors are conserved across species, we compared the behaviors of *S. stercoralis* to those of the rat-infective nematode *Strongyloides ratti*. We found that like *S. stercoralis* iL3s, *S. ratti* iL3s repeatedly push and puncture rat skin until penetration is complete (Fig. 2A, Movie S2). Interestingly, *S. ratti* iL3s exhibited more pushes, and began pushing earlier, on rat skin than *S. stercoralis* iL3s (Fig. 2B-D). These results suggest that although the two *Strongyloides* species engage in similar behaviors on skin, sensory cues specific to host skin increase the frequency of these behaviors. To test whether *S. stercoralis* also modulates its skin-penetration behavior on host skin, we examined the behavior of *S. stercoralis* iL3s on human skin; skin samples were obtained either from the forearm of cadavers or from the abdomen or breast of surgical patients following plastic surgery (Fig. S2). We found that *S. stercoralis* iL3s pushed down on human skin more frequently than rat skin (Fig. 2E-F). In some cases, *S. stercoralis* iL3s paused locomotion and pushed continuously on human skin for 10 s or more (Fig. 2E); in contrast, we never detected prolonged bouts of pushing on rat skin. *S. stercoralis* iL3s also pushed down more quickly after placement on human skin than rat skin (Fig. 2G). However, *S. stercoralis* iL3s were equally able to penetrate human and rat skin (Fig. 2H), indicating that the increase in skin-penetration behaviors on human skin reflects a higher penetration drive rather than an increased ability to penetrate. The higher penetration drive on host skin might reflect an excitatory behavioral state that is triggered by host-specific chemosensory cues.

**Figure 2.**
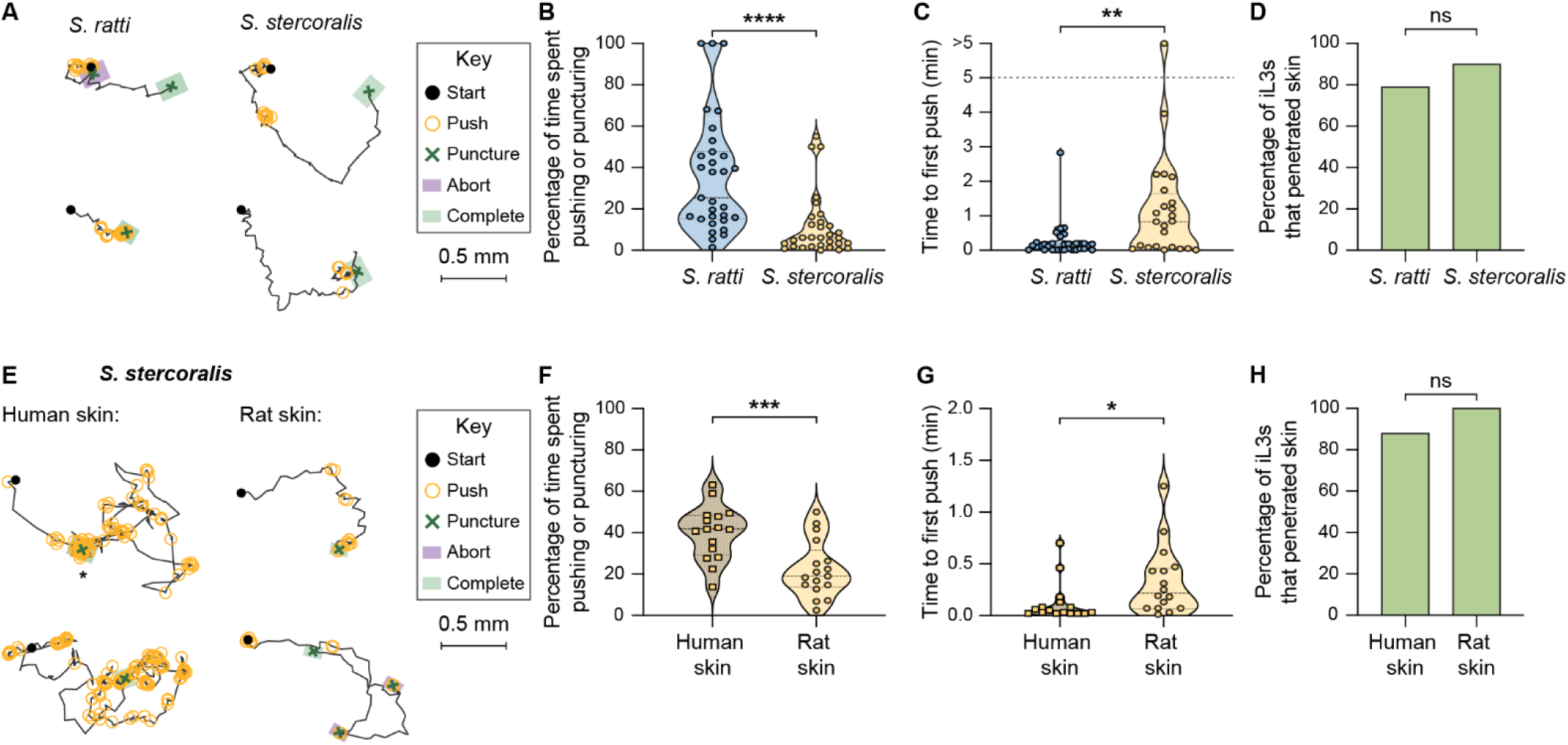
Skin-penetration behaviors are conserved between distinct species of skin-penetrating nematodes, but penetration drive is greater on host skin. **A.** *S. ratti* iL3s show similar skin-penetration behaviors to *S. stercoralis* iL3s. Tracks show the trajectories of two representative *S. ratti* iL3s and two representative *S. stercoralis* iL3s during skin penetration; the locations where one of four distinct behavioral motifs (push, puncture, aborted penetration, completed penetration) occurred along the worm paths are indicated by icons. During a push (yellow circle), the worm pushes down on the surface of the skin with its head; during a puncture (green X), the worm pushes its head into the skin; during an aborted penetration attempt (purple square), the worm stops attempting to enter skin and returns to the surface; and during a completed penetration attempt (green square), the worm fully enters the skin. The black lines indicate spaces where the worm is crawling on the surface of the skin in between attempts at penetration. A representative worm was one whose time spent pushing and puncturing skin and time to first puncture were close to the median value of the entire cohort that was tracked. **B.** *S. ratti* iL3s engage in more penetration attempts on rat skin than *S. stercoralis* iL3s. Violin plot shows the percentage of time on the skin surface that *S. ratti* and *S. stercoralis* iL3s spend engaging in pushes or punctures. n = 31 iL3s per species. \*\*\*\**p*<0.0001, Mann-Whitney test. The iL3s that had initiated penetration by the time the recording started were excluded from this analysis. **C.** *S. ratti* iL3s push down on rat skin more quickly than *S. stercoralis* iL3s. Violin plot shows the time between placement of iL3s on the skin surface and the first push event. n = 32-34 iL3s per species. \*\**p*<0.01, Mann-Whitney test. The iL3s that initiated penetration without detectable pushing were excluded from this analysis; pushing was likely not detected in these cases because the behavior occurred either before the recording started or too briefly to be captured at 2 frames/s. The dotted line at y = 5 indicates the time at which the assay ended; the dot above this line indicates a worm that failed to push on the skin by the end of the assay period. **D.** *S. ratti* and *S. stercoralis* iL3s penetrate rat skin with similar frequency. Bar graph shows the percentage of *S. ratti* and *S. stercoralis* iL3s that completed skin penetration. n = 31-34 iL3s per species. ns = not significant, Fisher’s exact test. **E.** *S. stercoralis* iL3s have higher penetration drive on human skin than rat skin. Two representative iL3s that were placed on either human skin or rat skin are shown; a representative worm was one whose time spent pushing and puncturing skin was close to the median value of the entire cohort that was tracked. The key shows behavioral motifs that were tracked. Asterisk indicates an instance when the iL3 pushed in one location for over 10 s; such prolonged pushing was never observed among iL3s on rat skin. **F.** *S. stercoralis* iL3s engage in more penetration attempts on human skin than rat skin. Violin plot shows the percentage of time that *S. stercoralis* iL3s spend pushing or puncturing the surface of human or rat skin. n = 16-17 iL3s per skin type. \*\*\**p*<0.001, unpaired t-test. **G.** *S. stercoralis* iL3s push down on human skin more quickly than rat skin. Violin plot shows the time taken by *S. stercoralis* iL3s to push down for the first time since placement on either human or rat skin. n = 16 iL3s per skin type. \**p*<0.05, Mann-Whitney test. **H.** *S. stercoralis* iL3s are equally able to penetrate human skin and rat skin. Bar graph shows the percentage of *S. stercoralis* iL3s that completed penetration on human or rat skin. n = 17 iL3s per species. ns = not significant, Fisher’s exact test. For B-C and F-G, dots depict individual worms, dashed lines indicate the median, and dotted lines indicate the interquartile range. Behavioral parameters plotted in B-D and F-H were each obtained from 4 independent replicate experiments. Skin from three distinct human donors was tested in the human skin assays.

### Pharmacological inhibition of dopamine receptors inhibits skin penetration in *Strongyloides* species and human hookworms

We next investigated the neural basis of skin-penetration behavior. Our behavioral analysis suggested that iL3s survey the texture of the skin surface in search of favorable entry points. In the free-living nematode *Caenorhabditis elegans*, the dopaminergic neurons mediate detection of textured surfaces^18-21^. We therefore hypothesized that dopamine signaling might regulate the detection of skin surface texture and drive the ensuing behavioral response in *S. stercoralis.* To test this, we treated *S. stercoralis* iL3s with haloperidol, which interferes with the activity of dopamine receptors^22,23^, and performed single-worm skin penetration assays. We found that haloperidol-treated iL3s crawled on the skin in circuitous paths, rarely pushed or punctured the skin, and often failed to penetrate (Fig. 3A-D). These behavioral phenotypes were rescued by the addition of exogenous dopamine (Fig. 3A-D), suggesting that haloperidol acts on the dopaminergic pathway to modulate skin penetration. Similar results were observed upon treatment of *S. ratti* iL3s with haloperidol and dopamine (Fig. S3). These results suggest a critical role for dopamine signaling in regulating skin penetration.

**Figure 3.**
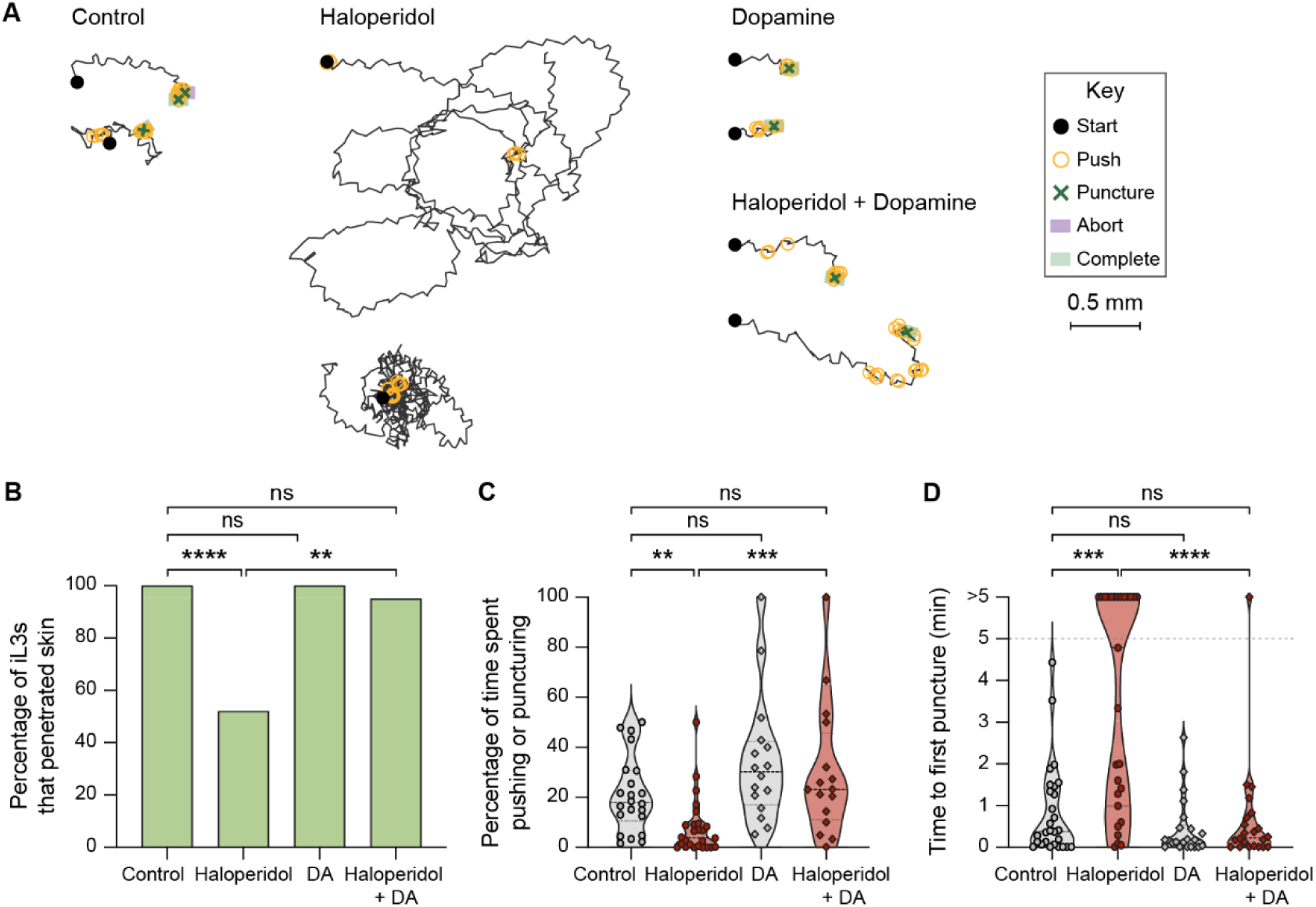
Pharmacological inhibition of dopamine signaling blocks skin penetration in *S. stercoralis*. **A.** Haloperidol inhibits skin penetration, and dopamine rescues this phenotype. Tracks show two representative *S. stercoralis* iL3s from each treatment group; for the haloperidol-treated group, two representative worms that did not puncture or complete penetration are shown. Representative worms that punctured and completed skin penetration were defined as those whose time to first puncture and time spent pushing and puncturing were close to the median value of the entire cohort that was tracked. Representative worms that neither punctured nor completed penetration were defined as those whose time spent pushing and puncturing the skin were close to the median value of the entire cohort. The key shows the behavioral motifs that were tracked. **B.** Haloperidol treatment of *S. stercoralis* iL3s inhibits skin penetration, and exogenous dopamine (DA) rescues this behavioral phenotype. Graph shows the percentage of iL3s treated with either the vehicle (control), 1.5 mM haloperidol, 10 mM DA, or 1.5 mM haloperidol + 10 mM DA that completed penetration. n = 22-28 iL3s per condition. \*\*\*\**p*<0.0001, \*\**p*<0.01, ns = not significant, Fisher’s exact test with a Bonferroni correction for multiple comparisons. **C.** Haloperidol reduces the percentage of time that *S. stercoralis* iL3s spend pushing and puncturing the skin, and this effect is rescued by addition of exogenous DA. Violin plot depicts the percentage of time that worms from each treatment group spent engaging in pushes and punctures. n = 16-26 iL3s per condition. \*\*\**p*<0.001, \*\**p*<0.01, ns = not significant, Kruskal-Wallis test with Dunn’s post-test. The iL3s that had initiated penetration by the time the recording started were excluded from this analysis. **D.** Haloperidol inhibits punctures, and DA rescues this behavioral phenotype. Violin plot depicts the time taken by worms from each treatment group to puncture the skin for the first time since placement on skin. The dotted line at y = 5 indicates the time at which the assay ended; the dots above this line indicate animals that failed to puncture the skin by the end of the assay period. n = 22-28 iL3s per condition. \*\*\*\**p*<0.0001, \*\*\**p*<0.001, ns = not significant, Kruskal-Wallis test with Dunn’s post-test. For C-D, dots depict individual worms, dashed lines indicate medians, and dotted lines indicate interquartile ranges. Behavioral parameters plotted in B-D were obtained from 4 independent replicate experiments.

To test whether dopamine signaling also regulates skin penetration in other human-infective nematodes, we repeated these experiments with the distantly related human-parasitic hookworm *Ancylostoma ceylanicum*. We found that treatment of *A. ceylanicum* iL3s with haloperidol also reduced pushes and punctures and inhibited skin penetration (Fig. 4A-C). Haloperidol-treated *A. ceylanicum* iL3s that failed to penetrate the skin also often failed to puncture the skin (Fig. 4D), showing that blocking dopamine signaling reduces the drive to penetrate. Dopamine rescued these behavioral defects (Fig. 4A-D). Together, our results demonstrate that dopamine signaling plays a conserved role in driving skin penetration across distantly related nematode species and highlight the potential for drugs that target the dopaminergic pathway to serve as broad-spectrum anti-nematode prophylactics.

**Figure 4.**
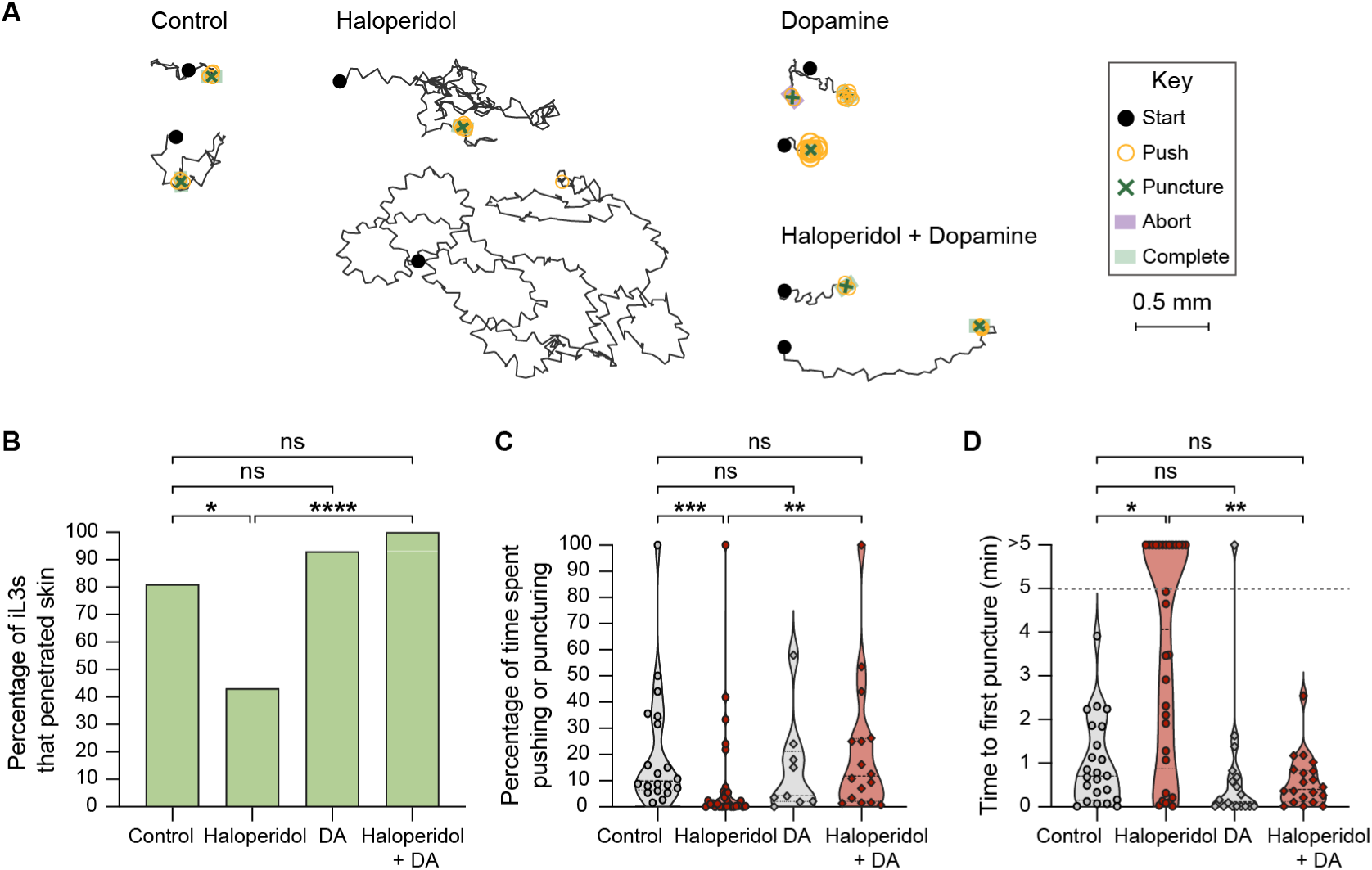
Pharmacological inhibition of dopamine signaling blocks skin penetration in the human-parasitic hookworm *A. ceylanicum*. **A.** Haloperidol inhibits skin penetration, and dopamine rescues this phenotype. Tracks show two representative worms from each treatment group; for the haloperidol-treated group, one representative worm that completed penetration and one randomly selected worm that did not complete penetration are shown. Representative worms were defined as in Fig. 3A. The key shows the behavioral motifs that were tracked. **B.** Haloperidol treatment of *A. ceylanicum* iL3s inhibits skin penetration and exogenous dopamine (DA) rescues this behavioral phenotype. Bar graph shows the percentage of iL3s that were treated with either the vehicle (control), 160 µM haloperidol, 10 mM DA, or 160 µM haloperidol + 10 mM DA that completed penetration. n = 14-30 iL3s per condition. \*\*\*\**p*<0.0001, \**p*<0.05, ns = not significant, Fisher’s exact test with a Bonferroni correction for multiple comparisons. **C.** Haloperidol reduces the percentage of time that *A. ceylanicum* iL3s spend pushing and puncturing the skin, and this effect is rescued by addition of exogenous DA. Violin plot depicts the percentage of time that worms from each treatment group spent engaging in pushes or punctures. n = 9-29 iL3s per condition. \*\*\**p*<0.001, \*\**p*<0.01, ns = not significant, Kruskal-Wallis test with Dunn’s post-test. iL3s that had initiated penetration by the time the recording started were excluded from this analysis. **D.** Haloperidol inhibits punctures and DA rescues this behavioral phenotype. Violin plot depicts the time taken by worms from each treatment group to puncture the skin since placement on skin. The dotted line at y = 5 indicates the time at which the assay ended; the dots above this line indicate worms that failed to puncture the skin by the end of the assay period. n = 18-30 iL3s per condition. \*\**p*<0.01, \**p*<0.05, ns = not significant, Kruskal-Wallis test with Dunn’s post-test. For C-D, dots depict individual worms, dashed lines indicate medians, and dotted lines indicate interquartile ranges. Behavioral parameters plotted in B-D were obtained from 4 independent replicate experiments.

### Disrupting dopamine biosynthesis blocks skin penetration

To directly test the role of dopamine signaling during skin penetration, we disrupted dopamine biosynthesis using CRISPR/Cas9-mediated targeted mutagenesis. We first mined the *S. stercoralis* genome and identified a putative homolog of the *C. elegans* gene *Ce-cat-2*, which encodes a tyrosine hydroxylase that mediates dopamine biosynthesis^24,25^ (Fig. 5A). In parallel, we also identified a putative homolog of the gene encoding the *C. elegans* dopamine transporter, *Ce-dat-1*^26^ (Fig. S4A). The amino acid sequences of *Ss-*CAT-2 and *Ce*-CAT-2 were 44% identical overall and 56% identical in the predicted catalytic domain^27^ (Fig. S4B). Similarly, *Ss-*DAT-1 and *Ce-*DAT-1 were 59.5% identical overall and 69.4% identical in the predicted monoamine transporter domain^27^(Fig. S4C). Transcriptional reporters for *Ss-cat-2* and *Ss-dat-1* were co-expressed in several cells that occupy the same position in iL3s as the *C. elegans* dopaminergic neurons^28,29^ (Fig. 5B). Specifically, the transcriptional reporters were expressed in a set of cells that send processes to the tip of the nose, which are likely the *Ss-*CEP neurons^24^; a cell immediately posterior to the candidate CEP neurons that is likely one of the two putative *Ss-* ADE neurons; and a cell along the body of the iL3 that is likely one of the two *Ss-*PDE neurons^24^. Together, the phylogenetic analysis and spatial expression profile of *Ss-cat-2* suggest that it is indeed the *S. stercoralis* tyrosine hydroxylase. Both *Ss-cat-2* and *Ss-dat-1* are significantly upregulated in iL3s relative to other life stages (Fig. 5C-D)^30,31^, consistent with a critical role for dopamine signaling in mediating iL3-specific behaviors.

**Figure 5.**
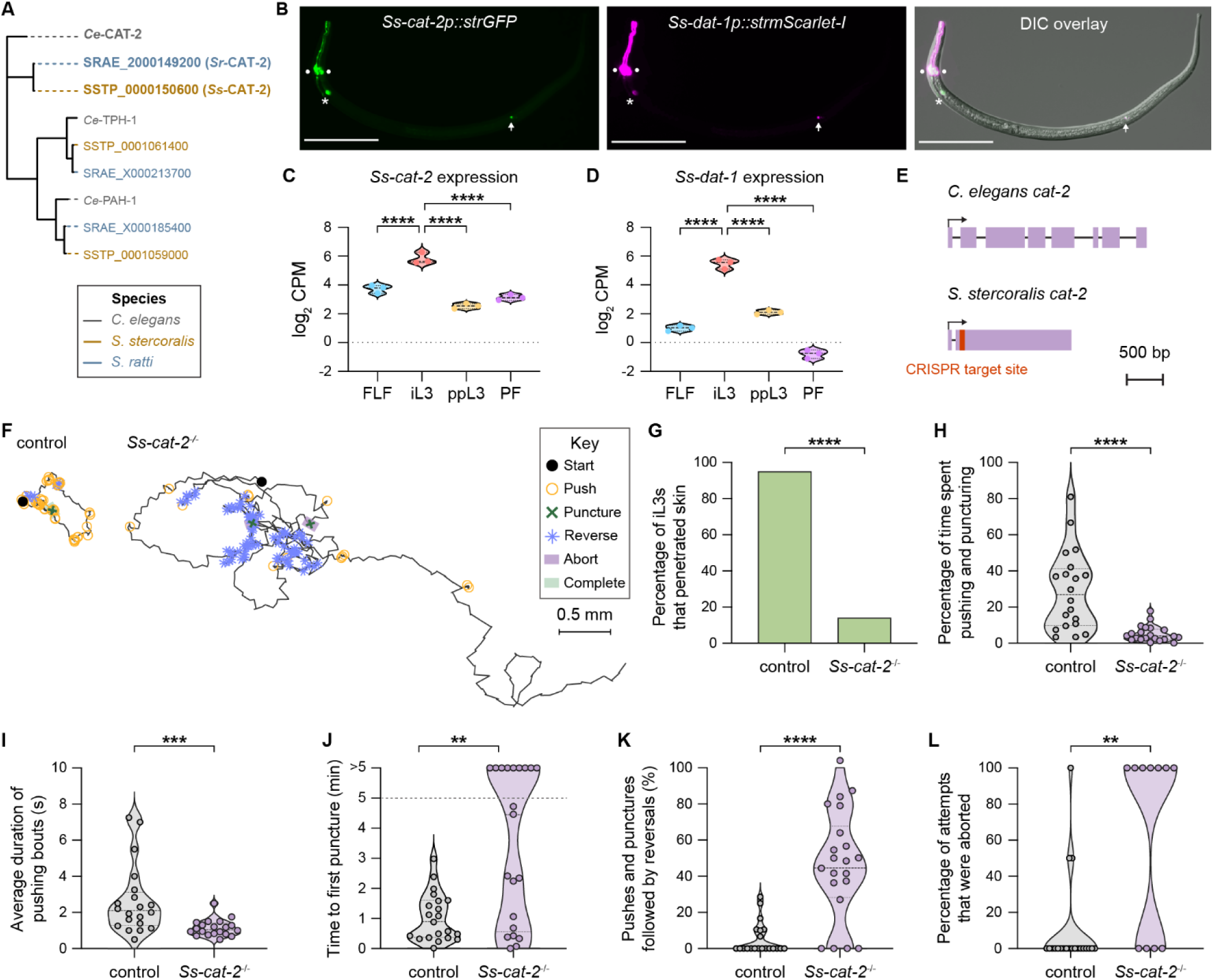
Dopamine is required for skin penetration. **A.** Phylogenetic analysis shows the closest homologs of *C. elegans* CAT-2 (gray) in *S. stercoralis* (brown) and *S. ratti* (blue). Putative homologs in each genome were identified by performing TBLASTN searches of the *C. elegans* CAT-2A protein sequence *(i.e*., the longest isoform) against either the *S. stercoralis* genome or the *S. ratti* genome in WormBase ParaSite WBPS18. The tree has the three members of the biopterin-dependent aromatic amino acid hydroxylases in *C. elegans*, of which CAT-2 is a member, and the predicted homologs in *S. stercoralis* and *S. ratti*. **B.** Co-expression of *Ss-cat-2* and *Ss-dat-1* in the putative dopaminergic (DA) neurons of *S. stercoralis*. Montage shows expression of the *Ss-cat-2* transcriptional reporter in green, expression of the *Ss-dat-1* transcriptional reporter in magenta, and co-expression of the two reporters and the relative positions of these neurons along the body of the iL3 in the differential interference contrast (DIC) overlay. The circles, asterisk, and arrow label the putative *Ss-*CEP, *Ss-* ADE, and *Ss-*PDE neurons, respectively. The iL3 is oriented with the dorsal side facing up and the ventral side facing down; the head is to the left. Scale bar = 100 µm. **C-D.** Violin plots show expression levels of *Ss-cat-2* and *Ss-dat-1*, expressed as log_2_ counts per million (CPM), at the indicated life stages based on published RNA-seq datasets^30,31,50^. \*\*\*\**p*<0.0001; statistical tests for differential expression analysis were performed as previously described, with corrections for multiple comparisons^30^. FLF = free-living female; iL3 = infective third-stage larva; ppL3 = post-parasitic third-stage larva; PF = parasitic female. Each dot indicates an independent replicate experiment. **E.** The *cat-2* genes of *C. elegans* and *S. stercoralis*. Schematics show the gene models of *Ce-cat-2* (isoform a), as annotated in WormBase WS292, and *Ss-cat-2*. The gene model for *Ss-cat-2* was initially derived from WBPS18 and then manually updated to include an additional exon^45^; the existence of this exon was supported by RNA-seq data^31,50^. Exons and introns are depicted as lavender boxes and black lines, respectively. The transcriptional start sites are indicated by black arrows. *Ss-cat-2* has a single CRISPR/Cas9 target site, which is located in the second exon (depicted in red). Drawings are to scale and scale bar = 500 bp. **F.** Inactivation of *Ss-cat-2* drastically alters skin-penetration behavior. Tracks show the skin-penetration behaviors of a representative wild-type iL3 that punctured and penetrated the skin and a representative *Ss-cat-2*^-/-^ iL3 that punctured but did not complete penetration. Representative worms were defined as in Fig. 3A. The key details the behavioral motifs that were tracked. **G.** Inactivation of *Ss-cat-2* severely impairs skin penetration. Bar graphs show the percentage of wild-type and *Ss-cat-2*^-/-^ iL3s that completed skin penetration. n = 21 iL3s per genotype. \*\*\*\**p*<0.0001, Fisher’s exact test. **H.** *Ss-cat-2^-/-^*iL3s pushed and punctured the skin for less time than control iL3s. Violin plot depicts the percentage of time on skin that control and *Ss-cat-2^-/-^* iL3s spent engaging in pushes or punctures. n = 20-21 iL3s per genotype. \*\*\*\**p*<0.0001, Mann-Whitney test. The iL3s that had initiated penetration by the time the recording started were excluded from this analysis. **I.** The pushing bouts of *Ss-cat-2^-/-^* iL3s are shorter than those of control iL3s. For each worm, the duration of each individual pushing bout was averaged and then plotted. n = 20 iL3s per genotype. \*\*\**p*<0.001, Mann-Whitney test. **J.** Inactivation of *Ss-cat-2* inhibits punctures. Violin plot depicts the time taken by control vs. *Ss-cat-2^-/-^*iL3s to puncture the skin for the first time since placement on skin. n = 21 iL3s per genotype. \*\**p*<0.01, Mann-Whitney test. The dotted line at y = 5 indicates the time at which the assay ended; the dots above this line indicate animals that failed to puncture the skin by the end of the assay period. **K.** *Ss-cat-2^-/-^* iL3s frequently reverse after a push or puncture event. Violin plot shows the percentage of pushes or punctures that were followed by backward locomotion that lasted at least 1 s for each genotype. n = 21 iL3s per genotype. \*\*\*\**p*<0.0001, Mann-Whitney test. **L.** *Ss-cat-2^-/-^* iL3s frequently abort penetration attempts. Violin plot depicts the percentage of penetration attempts, as defined by instances that the worm has punctured and partially entered the skin, that were aborted. n = 11-21 iL3s per genotype. \*\**p*<0.01, Mann-Whitney test. For H-L, dots depict individual worms, dashed lines indicate medians, and dotted lines indicate interquartile ranges. Behavioral parameters plotted in G-L were obtained from 3 independent replicate experiments.

We next disrupted the *Ss-cat-2* gene using CRISPR and then generated a stable mutant line by propagation of homozygous mutants in Mongolian gerbils, the laboratory host for *S. stercoralis*, as previously described^32^ (Fig. 5E, Fig. S4D-G). We found that inactivation of *Ss-cat-2* nearly eliminated skin-penetration behavior – ∼90% of the *Ss-cat-2^-/-^* mutants failed to penetrate both rat and human skin (Fig. 5F-G, Fig. S5A-B). Instead, *Ss-cat-2^-/-^*iL3s crawled in long, circuitous paths on the skin surface and rarely engaged in pushes (Fig. 5F, H, Fig. S5A, C). When they did push on the skin, the pushing bouts were ∼2-fold shorter in *Ss-cat-2^-/-^*iL3s relative to wild-type iL3s (Fig. 5I, Fig. S5D). Thus, inactivation of *Ss-cat-2* reduces the propensity of iL3s to halt locomotion and engage in focused pushing bouts. Nearly half of the *Ss-cat-2^-/-^* iL3s failed to puncture rat skin (Fig. 5J), and two-thirds failed to puncture human skin (Fig. S5E). Interestingly, *Ss-cat-2^-/-^* iL3s also frequently reversed after a push or puncture, whereas wild-type iL3s did not (Fig. 5K, Fig. S5F); this likely explains why the majority of the mutants that did puncture the skin subsequently backed out of the skin and aborted the penetration attempt (Fig. 5L). Together, our results indicate that dopamine signaling controls multiple facets of skin-penetration behavior: pushes, punctures, and the ability to successfully complete penetration following a puncture.

### Dopaminergic neurons drive skin penetration

To further examine the role of dopamine signaling in mediating skin penetration, we chemogenetically silenced the dopaminergic neurons using the histamine-gated chloride channel HisCl1^33^ (Fig. 6A, Fig. S6A). We found that like *Ss-cat-2*^-/-^ iL3s, iL3s with silenced dopaminergic neurons rarely engaged in pushes and punctures and instead actively crawled across the skin surface (Fig. 6B-D). More than half of the iL3s failed to penetrate the skin (Fig.6C) and this was often because of a failure to puncture the skin (Fig. 6E). These results indicate that the dopaminergic neurons of *S. stercoralis* drive skin-penetration behaviors. Histamine treatment of wild-type iL3s that did not express the *HisCl1* transgene did not significantly alter skin-penetration behavior, indicating that the behavioral phenotypes described above are specific to silencing of the dopaminergic neurons (Fig. S6B-D). We note that HisCl1-mediated silencing may not result in complete loss of dopaminergic neuron activity, which is likely why the phenotype of the iL3s with silenced dopaminergic neurons was slightly less severe than that of the *Ss-cat-2*^-/-^ iL3s.

**Figure 6.**
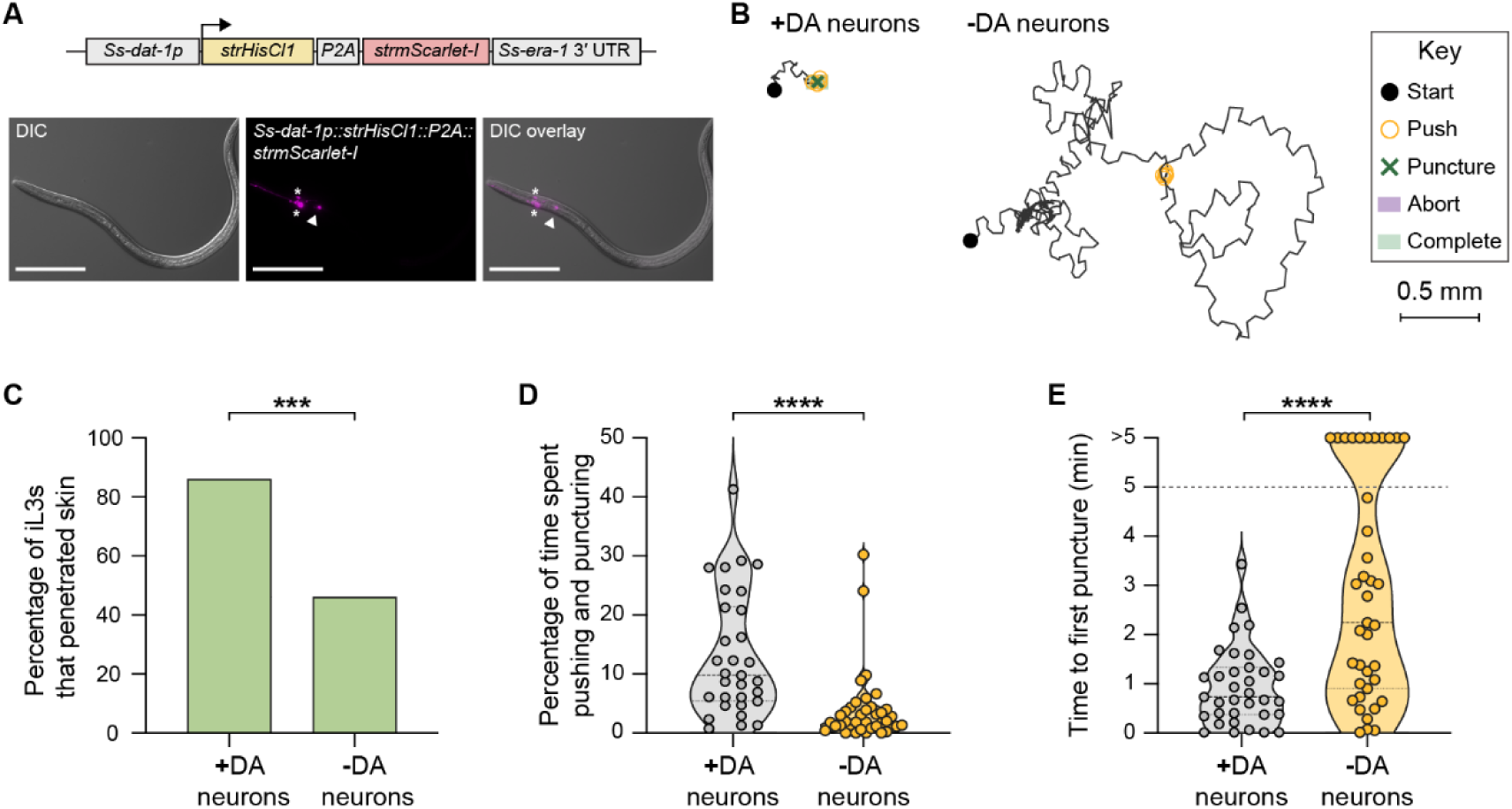
Dopaminergic neurons are required for skin penetration. **A.** Schematic shows the transgene used for chemogenetic silencing of the dopaminergic (DA) neurons of *S. stercoralis*. The histamine-gated chloride channel HisCl1 was expressed in the DA neurons using the promoter of the *Ss-dat-1* gene. Exposure of the transgenic *S. stercoralis* iL3s to exogenous histamine partially or fully silences the DA neurons (-DA neurons) relative to the vehicle-only control (+DA neurons). *Ss-dat-1p* = the *S. stercoralis dat-1* promoter; *strHisCl1* = *Strongyloides*-codon-optimized *HisCl1* gene; *P2A* = sequence encoding the self-cleaving P2A peptide; *strmScarlet-I* = *Strongyloides*-codon-optimized *mScarlet-I* gene; *Ss-era-1* 3′ UTR = the 3′ UTR of the *Ss-era-1* gene. Montage shows a representative transgenic iL3 that expresses the bicistronic *Ss-dat-1p::strHisCl1::P2A::strmScarlet-I* construct in the putative *Ss-*CEP neurons (asterisks) and *Ss-*ADE neurons (arrowhead). Fluorescence signal from the *Ss-*PDE neurons is rarely observed due to mosaic expression of this construct; thus, these neurons may not be silenced in our assays. Worm is oriented with the ventral side facing up and the head to the left. Scale bar = 100 µm. **B.** Silencing the DA neurons inhibits skin-penetration behavior. Behaviors of a representative mock-treated iL3 (+DA neurons) that punctured and completed penetration and a representative histamine-treated iL3 (-DA neurons) that neither punctured nor completed penetration are shown. Representative worms were defined as in Fig. 3A. The behavioral motifs that were tracked are detailed in the key. **C.** Fewer iL3s penetrate skin when DA neurons are silenced. Bar graph shows the percentage of mock-treated (+DA neurons) and histamine-treated (-DA neurons) iL3s that completed skin penetration. n = 35-39 iL3s per condition. \*\*\**p*<0.001, Fisher’s exact test. **D.** Silencing the DA neurons reduces pushes and punctures. Violin plot depicts the percentage of time on skin that mock-treated (+DA neurons) and histamine-treated (-DA neurons) iL3s spent engaging in pushes or punctures. n = 31-38 iL3s per condition. \*\*\*\**p*<0.0001, Mann-Whitney test. The iL3s that had initiated penetration by the time the recording started were excluded from this analysis. **E.** Silencing the DA neurons inhibits punctures. Violin plot depicts the time taken by mock-treated (+DA neurons) and histamine-treated (-DA neurons) iL3s to puncture the skin for the first time since placement on skin. n = 35-39 iL3s per condition. \*\*\*\**p*<0.0001, Mann-Whitney test. The dotted line at y = 5 indicates the time at which the assay ended; the dots above this line indicate animals that failed to puncture the skin by the end of the assay period. For D-E, dots depict individual worms, dashed lines indicate medians, and dotted lines indicate interquartile ranges. Behavioral parameters plotted in C-E were obtained from 6 independent replicate experiments.

### *Ss*-TRP-4 represents a possible target for topical prophylactic intervention

Are there nematode-specific components of the dopaminergic pathway that could be targeted for nematode control without interfering with host dopamine signaling? We hypothesized that the transient receptor potential channel TRP-4 might be one such target. In *C. elegans*, *Ce*-TRP-4 is expressed in the dopaminergic neurons and couples mechanosensation of textured surfaces (*e.g.*, bacterial lawns) with behavioral responses^19,20,34^. We therefore asked whether TRP-4 is conserved in *S. stercoralis* and if so, whether it might have been co-opted in the *S. stercoralis* dopaminergic neurons to drive skin-penetration behaviors. We identified a one-to-one homolog of *Ce*-TRP-4 in the *S. stercoralis* genome (Fig. 7A); the amino acid sequences of *Ss*-TRP-4 and *Ce*-TRP-4 were 57.1% identical (Fig. S7A). An *Ss-trp-4* transcriptional reporter was co-expressed with *Ss-dat-1* in the putative *Ss-*CEP and *Ss-*ADE neurons (Fig. 7B), and expression of the *Ss-trp-4* gene was significantly upregulated in the iL3 stage relative to other life stages (Fig. 7C)^30,31^. Taken together, these findings indicate that *Ss-trp-4* is expressed in the *S. stercoralis* dopaminergic neurons and might play an important role in iL3-specific behaviors.

**Figure 7.**
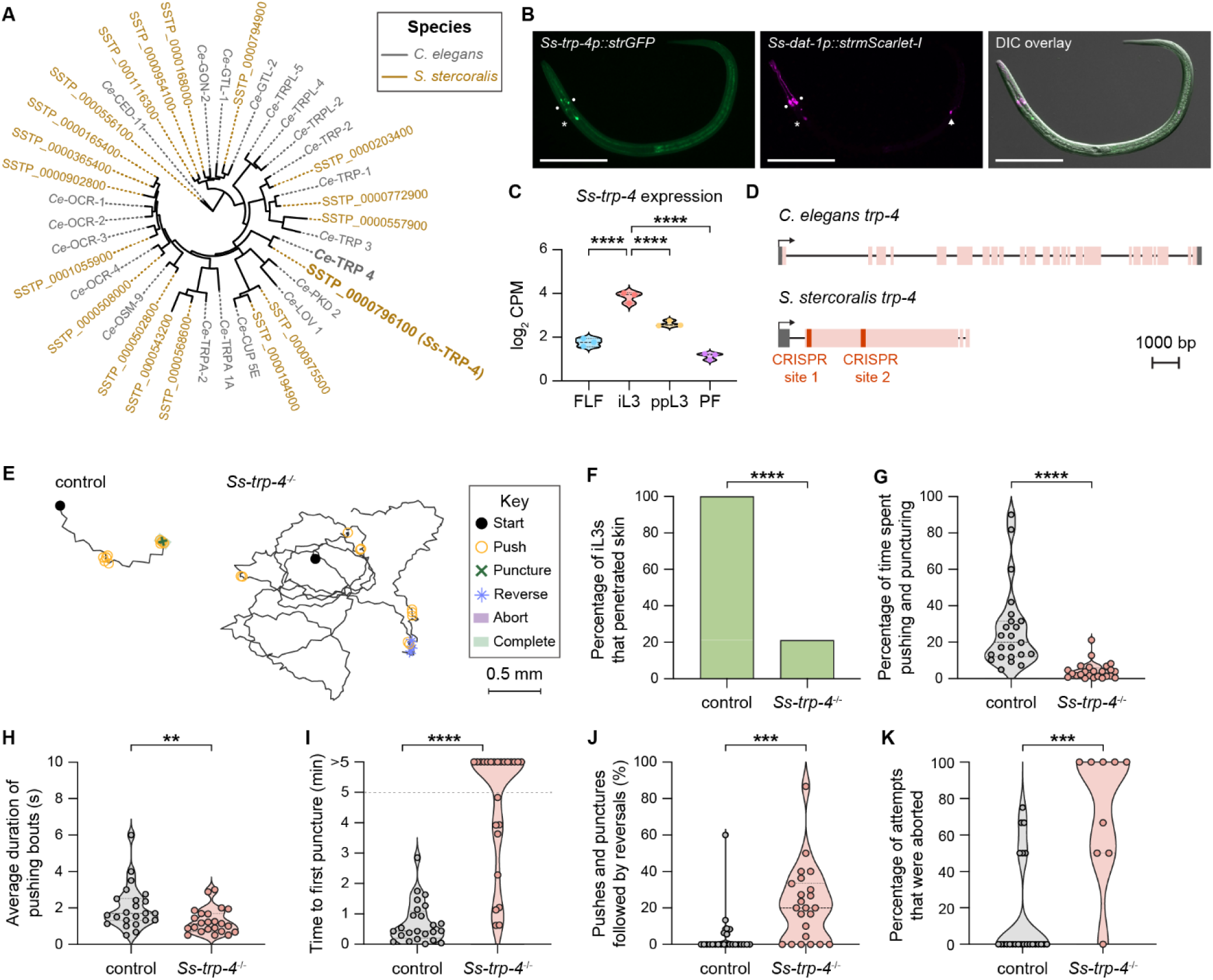
*Ss*-TRP-4 is required for skin penetration. **A.** Phylogenetic analysis shows the closest homologs of *C. elegans* TRP-4 (gray) in *S. stercoralis* (brown). Putative homologs in each genome were identified by performing TBLASTN searches of the *C. elegans* TRP-4 protein sequence against the *S. stercoralis* genome in WBPS18. The tree has all known TRP family members in *C. elegans*^51-53^ and the predicted homologs in *S. stercoralis*. **B.** Co-expression of *Ss-trp-4* and *Ss-dat-1* in the putative dopaminergic (DA) neurons of *S. stercoralis*. Montage shows expression of the *Ss-trp-4* transcriptional reporter in green, expression of the *Ss-dat-1* transcriptional reporter in magenta, and co-expression of the two reporters and the relative positions of these neurons along the body of the iL3 in the DIC overlay. The circles, asterisk, and arrow label the putative *Ss-*CEP, *Ss-*ADE, and *Ss-*PDE neurons, respectively; we never observed expression of the *Ss-trp-4* reporter in *Ss-*PDE. The iL3 is oriented with the dorsal side facing up and head to the left. Scale bar = 100 µm. **C.** Violin plot shows expression levels of *Ss-trp-4*, expressed as log_2_ counts per million (CPM), in the indicated life stages based on published RNA-seq datasets^30,31,50^. \*\*\*\**p*<0.0001; statistical tests for differential expression analysis were performed as previously described, with corrections for multiple comparisons^30^. FLF = free-living female; iL3 = infective third-stage larva; ppL3 = post-parasitic third-stage larva; PF = parasitic female. Each dot indicates an independent replicate experiment. **D.** The *trp-4* genes of *C. elegans* and *S. stercoralis*. Schematics show the gene models of *Ce-trp-4* and *Ss-trp-4*, which were derived from annotations in WS292 and WBPS18, respectively. Notably, the *Ss-trp-4* gene model has a 5′ UTR that is not annotated in WBPS18 but is supported by published RNA-seq data^31,50^. Exons, introns, and UTRs are depicted as pink boxes, black lines, and gray boxes, respectively. The transcriptional start sites are indicated by black arrows. *Ss-trp-4* has two CRISPR/Cas9 target sites in the first exon, which are depicted in red; we used two distinct sgRNAs, one targeting each CRISPR site, to inactivate *Ss-trp-4* and generate a mutant stable line. Drawings are to scale and scale bar = 1000 bp. **E.** *Ss-trp-4* mutants have reduced skin-penetration drive. Tracks show the skin-penetration behaviors of a representative wild-type iL3 that punctured and completed penetration and a representative *Ss-trp-4*^-/-^ iL3 that neither punctured nor completed penetration. Representative worms were defined as in Fig. 3A. The key details the behavioral motifs that were tracked. **F.** Inactivation of *Ss-trp-4* severely inhibits skin penetration. Bar graph shows the percentage of wild-type and *Ss-trp-4*^-/-^ iL3s that completed skin penetration. n = 24 iL3s per genotype. \*\*\*\**p*<0.0001, Fisher’s exact test. **G.** *Ss-trp-4^-/-^*iL3s pushed and punctured the skin for less time than control worms. Violin plot depicts the percentage of time on skin that control and *Ss-trp-4^-/-^*iL3s spent engaging in pushes or punctures. n = 23-24 iL3s per genotype. \*\*\*\**p*<0.0001, Mann-Whitney test. The iL3s that had initiated penetration by the time the recording started were excluded from this analysis. **H.** The pushing bouts of *Ss-trp-4^-/-^* iL3s are shorter than those of control iL3s. For each worm, the duration of each individual pushing bout was averaged and then plotted. n = 23 iL3s per genotype. \*\**p*<0.01, Mann-Whitney test. **I.** Inactivation of *Ss-trp-4* inhibits punctures. Violin plot depicts the time taken by control vs. *Ss-trp-4^-/-^* iL3s to puncture the skin for the first time since placement on skin. n = 24 iL3s per genotype. \*\*\*\**p*<0.0001, Mann-Whitney test. The dotted line at y = 5 indicates the time at which the assay ended; the dots above this line indicate animals that failed to puncture the skin by the end of the assay. **J.** *Ss-trp-4^-/-^* iL3s frequently reverse after a push or puncture event. Violin plot shows the percentage of pushes or punctures that were followed by backward locomotion that lasted at least 1 s for each genotype. n = 23-24 iL3s per genotype. \*\*\**p*<0.001, Mann-Whitney test. **K.** *Ss-trp-4^-/-^* iL3s frequently abort penetration attempts. Violin plot depicts the percentage of penetration attempts, as defined by instances that the worm has punctured and partially entered the skin, that were aborted. n = 9-24 iL3s per genotype. \*\*\**p*<0.001, Mann-Whitney test. For G-K, dots depict individual worms, dashed lines indicate the median, and dotted lines indicate the interquartile range. Behavioral parameters plotted in F-K were obtained from 3 independent replicate experiments.

To test whether *Ss-trp-4* is necessary for skin penetration, we generated a stable *Ss-trp-4^-/-^* knockout line^32^, as described above (Fig. 7D, Fig. S4F, and Fig. S7B-D). We then performed skin penetration assays with *Ss-trp-4^-/-^* and wild-type iL3s on both rat and human skin. Similar to inactivation of *Ss-cat-2*, inactivation of *Ss-trp-4* drastically reduced skin penetration (Fig. 7E-F, Fig. S8A-B). *Ss-trp-4^-/-^* iL3s pushed on the skin less frequently than control iL3s (Fig. 7E, G, Fig. S8C), and the pushing bouts that did occur were shorter and more often followed by a reversal (Fig. 7H, J, Fig. S8D). Moreover, over 60% of the mutants on rat skin and 50% of the mutants on human skin failed to puncture the skin at all (Fig. 7I, Fig. S8E), and the penetration attempts that were initiated were often aborted (Fig. 7K, Fig. S8F). These results show that *Ss*-TRP-4 plays a key role in driving skin penetration in *S. stercoralis*. Notably, *Ss-*TRP-4 is not conserved to humans^35^ (Fig. S9) but is conserved to the human-infective hookworm *A. ceylanicum* (Fig. S10). Thus, topical compounds containing drugs that target *Ss-*TRP-4 have the potential to prevent infections by multiple species of skin-penetrating nematodes while causing little to no side effects in humans.

## DISCUSSION

Here, we conduct an in-depth analysis of skin penetration, a critical but previously unstudied behavior that enables skin-penetrating nematodes to invade human hosts. We show that iL3s execute a complex set of behavioral motifs that allows them to probe the skin surface for favorable points of entry and culminates in the penetration of skin tissue (Fig. 1). Favorable entry points could include skin surface contours that enable iL3s to gain traction for burrowing into the skin, gaps between individual epidermal keratinocytes that provide reduced resistance to burrowing, hair follicles^4^, and open wounds. The finding that infective larvae actively explore the skin surface to locate favorable entry points, rather than entering the skin immediately upon contact, identifies a window of opportunity for preventative interventions and suggests that topical compounds that interfere with skin-penetration behaviors could be developed into the first anti-nematode prophylactic treatments.

We also uncover neural and molecular mechanisms that underlie skin penetration. By treating worms with haloperidol, a drug that interferes with the activity of the dopamine receptors, we show that dopamine signaling is necessary for skin penetration by *S. stercoralis*, *S. ratti,* and *A. ceylanicum* (Fig. 3-4 and Fig. S3). *S. stercoralis* and *A. ceylanicum* occupy distinct phylogenetic clades, both of which are thought to have evolved from free-living ancestors^36^. Thus, our findings imply that the role of dopamine signaling in driving skin penetration evolved independently in the two species of parasitic nematodes. Going further, we show that the dopamine biosynthesis enzyme *Ss-*CAT-2, the dopaminergic neurons, and the putative mechanoreceptor *Ss*-TRP-4 mediate skin-penetration behavior in *S. stercoralis*. *Ss*-TRP-4 is conserved to *A. ceylanicum*, but not to humans, suggesting that preventative interventions that target this protein may be both broadly effective against diverse species of skin-penetrating nematodes and safe for administration to humans.

Skin penetration is a parasite-specific behavior. Sensory neuroanatomy is largely conserved across nematode species^37,38^, raising the question of how parasite-specific behaviors have evolved in parasitic nematodes despite their having essentially the same sensory neuroanatomy as free-living nematodes. In *C. elegans*, the dopaminergic neurons are necessary for the worms to slow down upon entry into a patch of food, and treatment of *C. elegans* with exogenous dopamine halts locomotion^18,20,39^. In skin-penetrating nematodes, the dopaminergic neurons have been co-opted to drive skin penetration. Our data suggest that dopamine signaling causes the worm to stop crawling and instead push its head against the skin. Indeed, we show that the average duration of pushing bouts in the *Ss-cat-2^-/-^*mutants is shorter than wild type (Fig. 5I, Fig. S5D), indicating that *Ss-cat-2^-/-^*iL3s have a reduced propensity to halt locomotion and push down on skin. Thus, while dopamine signaling appears to cause pauses in locomotion in both free-living and skin-penetrating nematodes, in skin-penetrating nematodes the dopaminergic neurons are coupled to an additional downstream motor program that stimulates skin pushing and ultimately skin penetration.

How might dopamine signaling promote skin penetration? Our results suggest a model where *Ss*-TRP-4 acts as a mechanotransduction channel in the sensory endings of the dopaminergic neurons that opens when iL3s encounter potential entry points on host skin, causing excitation of the neurons. The dopaminergic neurons then release dopamine, which binds to downstream dopamine receptors, causing the worm to stop crawling and instead push against the skin. Continued pushing, coupled with the secretion of metalloproteases that help break down the skin^4^, leads to skin penetration. Our results also suggest that dopamine signaling suppresses mechanosensory behaviors that would otherwise prevent skin penetration. *Ss-cat-2^-/-^* iL3s often reversed after a push or puncture, whereas wild-type iL3s did not (Fig. 5K); these reversals prevented the forward locomotion required for skin penetration. The increased tendency of *Ss-cat-2^-/-^* iL3s to reverse after pushes and punctures might reflect a hypersensitivity of the mutants to nose touch, as both pushes and punctures are always preceded by head-on collisions with the skin surface. This model is consistent with studies that have shown that dopamine signaling modulates the sensitivity of *C. elegans* to stimuli that cause reversal behavior, including nose touch^23,40,41^. *Ss*-TRP-4 may mediate the effect of dopamine signaling on nose-touch sensitivity, as *Ss-trp-4* mutants also reverse frequently after a push or puncture (Fig. 7J). The dopamine signaling pathway could act constitutively or in a context-dependent manner (*e.g.*, when the worm is on skin) to modulate nose-touch sensitivity.

Can infective larvae distinguish host from non-host skin? Our comparison of the skin-penetration behaviors of *S. stercoralis* and *S. ratti* revealed that host selectivity is not regulated at the level of skin penetration ability, as ∼90% of the tested *S. stercoralis* iL3s penetrated rat skin. However, *S. stercoralis* iL3s pushed on human skin more frequently and earlier than rat skin (Fig. 2E-G), indicating that *S. stercoralis* iL3s can distinguish host from non-host skin and have a higher drive to penetrate human skin. Host-specific cues, such as skin odorants and metabolites, might evoke this increased penetration drive^41^.

In summary, our results illuminate the complex sequence of behaviors executed by skin-penetrating nematodes on skin and highlight the key role of dopamine signaling in driving these behaviors. Our study also demonstrates that exposing skin-penetrating nematodes to pharmacological inhibitors of dopamine signaling can block skin invasion. While topical insect repellents are widely employed to prevent the spread of insect-transmitted diseases, the possibility of developing topical repellents for parasitic nematodes has not been explored. Our results illustrate the potential for topically applied compounds that block TRP-4 or other nematode-specific components of the dopaminergic pathway to be developed into novel anti-nematode prophylactics.

## DATA AVAILABILITY

All data necessary for the conclusions described in this study are included with this article. Custom code used for skin penetration tracking assays is available from GitHub (https://github.com/BryantLabUW/WormTracker3000.git). Source data are provided with this paper (Dataset S1).

## METHODS

### Ethics statement

All animal protocols and procedures were approved by the UCLA Office of Animal Research Oversight (Protocol ARC-2011-060). The protocol follows the guidelines set by the AAALAC and the *Guide for the Care and Use of Laboratory Animals*. Human skin samples were collected following approval by the University of California Institutional Review Board (Protocol 22-000400), with signed written informed consent obtained in accordance with the Declaration of Helsinki principles.

### Strains

The following *S. stercoralis* strains were used in this study: UPD, EAH435 *bruIs4*[*Ss-act-2p::strmScarlet-I:: Ss-era-1* 3′ UTR]^15^, EAH477 *Ss-cat-2(bru3[Ss-act-2p::strmScarlet-I])* II */ Ss-cat-2(bru4[Ss-act-2p::strGFP])* II, and EAH489 *Ss-trp-4(bru5[Ss-act-2p::strmScarlet-I]*) II */ Ss-trp-4(bru6[Ss-act-2p::strElectra2::P2A::strElectra2])* II. The EAH477 strain is composed of worms that express both mScarlet-I and GFP, mScarlet-I only, and GFP only. The dual-colored worms have the *strmScarlet-I* transgene inserted in one allele of *Ss-cat-2* and the *strGFP* transgene inserted in the other allele; the single-colored worms have the same transgene inserted into both alleles of *Ss-cat-2*. In all cases, the transgenes are inserted into the CRISPR site 5′-GTATCTAATTTTGCTAATGG-3′ in *Ss-cat-2*. Similarly, EAH489 is composed of worms that express both mScarlet-I and Electra2, mScarlet-I only, and Electra2 only. The dual-colored worms have the *strmScarlet-I* transgene inserted in one allele of *Ss-trp-4* and the *strElectra2::P2A::strElectra2* transgene inserted in the other allele; the single-colored worms have the same transgene inserted into both alleles of *Ss-trp-4*. In all cases, the transgenes are inserted between the CRISPR sites 5′-GGCTCTTCAAGTAAACCAGG-3′ and 5′-GCTGATGTTCATTTACATGG-3′ in *Ss-trp-4*. The *S. ratti* ED231 strain and the *A. ceylanicum* Indian strain (US National Parasite Collection Number 102954) were also used in this study.

### Maintenance of Strongyloides stercoralis

*S. stercoralis* was serially passaged in Mongolian gerbils (Charles River Laboratories). Outside of the gerbil, *S. stercoralis* was maintained on fecal-charcoal plates, as previously described^42^. Gerbil infections were done by first collecting *S. stercoralis* iL3s from fecal-charcoal plates using a Baermann apparatus^43^ and then washing the iL3s 5 times in sterile 1X PBS. After the last wash, the worms were resuspended in 1X PBS at a concentration of ∼10 worms/µL. Each gerbil was anesthetized using isoflurane and then inoculated by subcutaneous injection of 200 µL of the worm/PBS suspension, resulting in an infective dose of ∼2,000 iL3s/gerbil; 8-12 gerbils at a time were used for strain maintenance. Feces were collected from days 14-44 post-inoculation by housing gerbils overnight on wire racks, over damp cardboard lining (Shepherd Techboard, 8 x 16.5 inches, Newco, 999589), in cages. Each morning, feces were collected from the cardboard and then mixed with dH_2_O and autoclaved charcoal granules (Bone char, 4 lb pail, 10 x 28 mesh, Ebonex). Fecal-charcoal mixtures were packed, on top of damp Whatman paper, into 10 cm Petri plates (VWR, 82050-918) and placed in plastic boxes lined with damp paper towels. These boxes were either placed directly in a 23°C incubator or kept at 20°C for two days and then moved to a 23°C incubator. *S. stercoralis* free-living females for microinjection were collected from fecal-charcoal plates kept either at 25°C for one day or 20°C for two days.

### Maintenance of Strongyloides ratti

*S. ratti* was serially passaged in Sprague-Dawley rats (Inotiv). Outside of the rat, *S. ratti* was maintained on fecal-charcoal plates. Rat infections were done by first collecting *S. ratti* iL3s from fecal-charcoal plates using a Baermann apparatus^43^ and then washing the iL3s 5 times in sterile 1X PBS. Each rat was anesthetized with isoflurane and infected, via subcutaneous injection, with ∼700 iL3s in 200 µL of sterile PBS; 2-4 rats at a time were used for strain maintenance. Feces were collected as described above for *S. stercoralis*, except that collections were done from days 7-21 post-inoculation. Fecal-charcoal mixtures were made and maintained as described above for *S. stercoralis*.

### Maintenance of Ancylostoma ceylanicum

*A. ceylanicum* was serially passaged in male Golden Syrian hamsters (Inotiv). Outside of hamsters, *A. ceylanicum* was maintained on fecal-charcoal plates. Each hamster was infected by oral gavage with 60-100 iL3s resuspended in 100 µL of sterile 1X PBS. Between 2-8 hamsters at a time were used for strain maintenance. Feces were collected and fecal-charcoal mixtures were made and maintained as described above for *S. stercoralis*.

### Phylogenetic analyses

We identified the putative *Strongyloides* homologs of *Ce*-DAT-1 using a previously described approach^44^. Briefly, the protein sequence of *Ce*-DAT-1 was retrieved from WormBase WS292 and all similar genes in the *S. stercoralis* and *S. ratti* genomes in WormBase Parasite WBPS18 were identified by performing TBLASTN searches. The accuracy of the gene models for the hits in the *S. stercoralis* and *S. ratti* genomes was checked as described^44,45^; any revisions to gene models were done manually in Geneious Prime 2022.2. The protein sequences of each of these genes were then used in reciprocal TBLASTN searches against WS292. This approach identified all members of the sodium neurotransmitter symporter family (SNF) of proteins in *C. elegans*, including *Ce*-DAT-1. A MUSCLE alignment of all the protein sequences retrieved from the three nematode genomes was performed in Geneious Prime 2022.2. The alignment was fed into IQ-TREE (version 1.6.12), which used a VT+I+G4 substitution model to generate a phylogenetic tree^46,47^; the tree was then visualized using the interactive Tree of Life (iTOL)^48^. Using this approach, we identified SSTP_0000953300 and SRAE_2000329000 as the *S. stercoralis* and *S. ratti* homologs of *Ce-*DAT-1, respectively. Additionally, the monoamine transporter domain was identified in both *Ce*-DAT-1 and *Ss*-DAT-1 by performing CD-Search^49^ with corresponding protein sequences in the Conserved Domain Database.

The same approach was used to identify the *Strongyloides* homologs of *Ce-*CAT-2. Here, the isoform A of *Ce-*CAT-2 was used in TBLASTN searches because it is the longest isoform. We identified SSTP_0000150600 and SRAE_2000149200 as the *S. stercoralis* and *S. ratti* homologs of *Ce-*CAT-2, respectively. The aromatic amino acid hydroxylase domain in *Ce*-CAT-2 and *Ss-*DAT-1 was identified using the CD-Search tool in the Conserved Domain Database^49^. Additionally, the version of SSTP_0000150600 annotated in WBPS18 was missing an upstream exon based on publicly available RNA-seq datasets^31,50^. This upstream exon was added manually to the front of the SSTP_0000150600 gene using Geneious Prime 2022.2.

This approach was also used to identify the *Strongyloides* homologs of *C. elegans* TRP channels^51-53^. The protein sequences of *C. elegans* TRP channels were retrieved from WS292 and used in TBLASTN searches. We identified SSTP_0000796100 and SRAE_200231000 as the *S. stercoralis* and *S. ratti* homologs of *Ce-*TRP-4, respectively. Additionally, the version of SSTP_0000796100 annotated in WBPS18 was missing a 5′ UTR based on publicly available RNA-seq datasets^31,50^. This 5′ UTR was added manually to the front of the SSTP_0000796100 gene using Geneious Prime 2022.2. While building the tree of the *C. elegans* and *S. stercoralis* TRP channels, we found a putative novel, longer isoform of the gene SSTP_0000543200, in which the third intron was not spliced, based on RNA-seq data^31,50^. Using the same RNA-Seq datasets^31,50^, we also found a putative upstream exon in the gene SSTP_0000194900. We manually revised these gene models in Geneious Prime 2022.2.

To build the phylogenetic tree with the TRP channels in *C. elegans*, *S. stercoralis*, *S. ratti*, humans, and mice, we first retrieved the protein sequences of known *H. sapiens* and *M. musculus* TRP family members^54^ from UniProtKB (accession numbers are listed in Table S2). A MUSCLE alignment of all the protein sequences was done in Geneious Prime 2022.2. As described above, the alignment was then fed into IQ-TREE (version 1.6.12) to generate a phylogenetic tree^46,47^, using the same substitution model as above. The tree was visualized using the interactive Tree of Life (iTOL)^48^.

We used the same approach as above to identify the *Ancylostoma ceylanicum* homologs of *C. elegans* TRP channels^51-53^. The protein sequences of *C. elegans* TRP channels were retrieved from WS292 and used in TBLASTN searches against the *A. ceylanicum* genome (PRJNA23179) in WormBase Parasite WBPS19. The protein sequences were aligned, and the phylogenetic tree was built and visualized as described above^46-48^.

### Molecular biology

The promoter of *Ss-dat-*1 in the plasmid pAGR02 (*Ss-dat-1p::strHisCl1::P2A::strmScarlet-I::Ss-era-1* 3′ UTR) was made by PCR-amplifying the 2,579 bp region immediately upstream of the start codon of *Ss-dat-1* (SSTP_0000953300, WBPS18) using the primers RP15 and RP16. The PCR product was then cloned upstream of the *strHisCl1::P2A::strmScarlet-I::Ss-era-1* 3′ UTR cassette, which is contained in pMLC207, using the restriction enzymes HindIII and AgeI. To generate pAGR04 (*Ss-dat-1p::strmScarlet-I::Ss-era-1* 3′ UTR), the *Ss-dat-1* promoter fragment from pAGR02 was excised using the restriction enzymes HindIII and AgeI and then cloned upstream of the *strmScarlet-I::Ss-era-1* 3′ UTR cassette, which is contained in the plasmid pMLC201, using these same restriction enzymes.

The promoter of *Ss-cat-2* in the plasmid pRP19 (*Ss-cat-2p::strGFP::Ss-era-1* 3′ UTR), corresponding to the region 1,639 bp upstream of the new start codon of *Ss-cat-2* (SSTP_0000150600, WBPS18) was synthesized by GenScript. GenScript then cloned this promoter fragment upstream of the *strGFP::Ss-era-1* 3′ UTR cassette, which is contained in pMLC200, using the restriction enzymes HindIII and AgeI to generate pRP19.

The promoter of *Ss-trp-4* in the plasmid pRP37 (*Ss-trp-4p::strGFP::Ss-era-1* 3′ UTR), corresponding to the region 2,320 bp upstream of the start codon of *Ss-trp-4* (SSTP_0000796100, WBPS18), was synthesized by GenScript. GenScript then cloned this promoter fragment upstream of the *strGFP::Ss-era-1* 3′ UTR cassette, which is contained in pMLC200, using the restriction enzymes EcoRV and AgeI to generate pRP37.

To generate pRP23 (*Sr-U6p::Ss-cat-2 sgRNA::sgRNA scaffold::Sr-U6* 3′ UTR), which is the plasmid that contains the CRISPR single guide RNA (sgRNA) for *Ss-cat-2*, we first used the Find CRISPR sites tool in Geneious Prime 2022.2 to search for guide RNA sites that matched the consensus sequence 5′-GN(17)GG-3′^55^. We found one such site in the second exon of *Ss-cat-2*, with the sequence 5′-GTATCTAATTTTGCTAATGG-3′ and a Doench (2016) activity score of 0.538^56^. This guide RNA sequence was synthesized and fused with the promoter of the *S. ratti* U6 gene, the sequence of the sgRNA scaffold, and the *S. ratti* U6 3′ UTR by GenScript, as previously described^55^. The plasmid pRP31 is the HDR cassette that was used to insert *Ss-act-2p::strmScarlet-I::Ss-era-1* 3′ UTR into the *Ss-cat-2* gene. To make this plasmid, a 479 bp fragment immediately upstream (5′ homology arm) and a 514 bp fragment immediately downstream (3′ homology arm) of the Cas9 cut site in *Ss-cat-2* were first synthesized by GenScript. The 5′ homology arm was cloned upstream of the *Ss-act-2p::strmScarlet-I::Ss-era-1* 3′ UTR cassette, which is contained in pRP12, using the enzymes HindIII and KpnI to generate pRP30. The 3′ homology arm was cloned downstream of the *Ss-act-2p::strmScarlet-I::Ss-era-1* 3′ UTR cassette in pRP30 using the enzymes EagI and BamHI to generate pRP31. The plasmid pRP44 contains the other HDR cassette, *Ss-act-2p::strGFP::Ss-era-1* 3′ UTR, which was made by replacing *strmScarlet-I* in pRP31 with *strGFP* from pMLC200 using the restriction enzymes AgeI and AvrII. Generation of the plasmid pPV540 (*Sr-eef1-1ap::strCas9::Ss-era-1* 3′ UTR) was previously described^55^.

To generate pRP41 (*Sr-U6p::Ss-trp-4 sgRNA1::sgRNA scaffold::Sr-U6* 3′ UTR) and pRP42 (*Sr-U6p::Ss-trp-4 sgRNA2::sgRNA scaffold::Sr-U6* 3′ UTR), which are the plasmids that contain CRISPR sgRNAs that target sites 1 and 2, respectively, in *Ss-trp-4* (Fig. 7D), we first used the Find CRISPR sites tool in Geneious Prime 2022.2, as described above, to identify CRISPR/Cas9 target sites. We identified site 1 (5′-GGCTCTTCAAGTAAACCAGG-3′) and site 2 (5′-GCTGATGTTCATTTACATGG-3′), which had Doench (2016) activity scores of 0.73 and 0.702^56^, respectively. The guide RNA sequences were synthesized and fused with the promoter of the *S. ratti* U6 gene, the sequence of the sgRNA scaffold, and the *S. ratti* U6 3′ UTR by GenScript, as previously described^55^. The plasmid pRP40 is the HDR cassette that was used to insert *Ss-act-2p::strmScarlet-I::Ss-era-1* 3′ UTR into the *Ss-trp-4* gene. To make this plasmid, a 511 bp fragment immediately upstream (5′ homology arm) and a 555 bp fragment immediately downstream (3′ homology arm) of CRISPR sites 1 and 2, respectively, in *Ss-trp-4* were first synthesized by GenScript. The 5′ homology arm was cloned upstream of the *Ss-act-2p::strmScarlet-I::Ss-era-1* 3′ UTR cassette, which is contained in pRP12, using the enzymes HindIII and KpnI to generate pRP38. The 3′ homology arm was cloned downstream of the *Ss-act-2p::strmScarlet-I::Ss-era-1* 3′ UTR cassette in pRP38 using the enzymes EagI and BamHI to generate pRP40. The plasmid pRP50 contains the other HDR cassette, *Ss-act-2p::strElectra2::P2A::strElectra2::Ss-era-1* 3′ UTR. To make this plasmid, the *Strongyloides* codon-optimized version of *Electra2*^57^ was first synthesized in the form of *strElectra2::P2A::strElectra2* by GenScript and inserted into the pRP31 backbone using the restriction enzymes AgeI and AvrII. This generated *Ss-act-2p::strElectra2::P2A::strElectra2::Ss-era-1* 3′ UTR. The 5′ homology arm for inactivation of *Ss-trp-4* was inserted upstream of the reporter cassette in pRP31 using the enzymes HindIII and KpnI to generate pRP49. The 3′ homology arm for inactivation of *Ss-trp-4* was inserted downstream of *Ss-act-2p::strElectra2::P2A::strElectra2::Ss-era-1* 3′ UTR in pRP49, using the restriction enzymes EagI and BamHI, to generate pRP50.

### Single-worm skin penetration tracking assays with *S. stercoralis* and *S. ratti* on rat skin

Skin (from the epidermis to the hypodermis) was retrieved from the dorsal and lateral sides of either male or female Sprague-Dawley rats that were 3-13 months old. Euthanized rats that were used for skin retrieval had been frozen at -80°C either 0 or 1 times; if they were previously frozen, they were allowed to thaw overnight, at room temperature, before the skin was harvested. After retrieval of the skin, it was sectioned into small pieces that were approximately 2 cm x 2 cm. The skin was then either frozen at -80°C or used immediately for skin penetration assays. For skin penetration assays, pieces of skin were first allowed to equilibrate to room temperature and fur was then manually plucked from the surface of the skin. Skin sections were then draped over plastic cell culture inserts (CellCrown) and placed in individual wells of either 12-well or 6-well plates. If a 12-well plate (VWR Scientific, 10062-894) was used, the skin was draped over cell culture inserts made for 24-well plates (Millipore Sigma, Z742381). If a 6-well plate (Corning, 3516) was used, the skin was placed on cell culture inserts made for 12-well plates (Millipore Sigma, Z742383). The wells were filled with 1-3 mL of BU saline^58^ prior to placing the skin and insert in the well so as to ensure moisture retention within the skin. For some assays, the skin was air-dried for 5-15 minutes before performing assays to remove excess moisture on the skin surface. If the skin was slightly dried for assays, both control and experimental groups were tested on skin that was dried to the same extent.

While the skin was equilibrating to room temperature, *S. stercoralis* or *S. ratti* iL3s were isolated from fecal-charcoal plates using a Baermann apparatus^43^, as previously described. The fecal-charcoal plates used for collection of *S. stercoralis* UPD, *S. ratti* ED231, *S. stercoralis* EAH435, *S. stercoralis* EAH477 and *S. stercoralis* EAH489 were 7-10 days old. The plates used for collection of transgenic *S. stercoralis* iL3s from free-living adults that were microinjected were 5-7 days old. If worms were stained with DiI prior to skin penetration assays, a ∼50-100 µL worm pellet from the Baermann apparatus was first resuspended in ∼10 mL of BU saline^58^ and this worm suspension was then used for dye staining. One mL of the suspension was dyed with 5 µL of DiI (2 mg/mL, Thermo Fisher Scientific, D3911) for 15 minutes prior to skin penetration assays; the solvent for DiI was N,N-dimethylformamide (DMF, Thermo Fisher Scientific, 68-12-2) and DiI solutions used for staining were no older than 3 months. After staining, the worms were washed twice in fresh BU^58^ to remove excess DiI and DMF from the surface of the worm. Next, the worms (either DiI-stained or unstained, transgenic) were plated onto 10 cm unseeded nematode growth medium (NGM) plates^59^. Individual worms were picked from NGM plates using a paintbrush and allowed to crawl onto the skin surface. Immediately thereafter, the worms on skin were recorded using a Leica M165 FC fluorescence dissection microscope, with the ET-mCherry filter set (Leica, 10450195), and an attached Basler ace (acA3800-14um) camera for either 5 or 10 min or until skin penetration was complete. A series of still images was captured at 4 frames/second (fps) and image acquisition was controlled with the pylon Viewer software (Basler). The field of view was manually adjusted whenever the worm moved out of the field of view of the microscope during the course of the assay. Up to 10 worms were assayed, sequentially, on the same piece of skin. Whenever wild-type and mutant worms were assayed on the same day, assays were performed blind to genotype and blinding was lifted after the experiment was over.

Images captured for each worm were opened in Fiji 2.9.0/1.53t^60^ using File>Import>Image Sequence. Although images were captured at 4 fps, every alternate image was opened in Fiji as this resolution was sufficient for tracking behavior. All behaviors that are described were scored manually. Worms normally pushed the skin and then either crawled on the surface or punctured the skin. A bout of pushing was defined as the first frame at which the worm was observed to be pushing on the skin until the frame just prior to the one in which the worm either was crawling or had punctured the skin. Worms were considered to be reversing if they moved backwards for 2 or more frames (*i.e.*, 1 s or more). A reversal was considered to be associated with a push or puncture if it occurred within 5 frames (*i.e.*, 2.5 s or less) of the latter behaviors. Consecutive frames of backward movement that lasted for at least 2 frames (*i.e.*, 1 s or more) and were interrupted for fewer than 4 frames were ascribed to the same reversal bout.

To generate tracks of worms on skin, worm position was tracked manually in Fiji using the TrackMate plugin^61^ and plotted using custom MATLAB software that can be accessed at this URL: https://github.com/BryantLabUW/WormTracker3000.git. In cases where the field of view was adjusted during the course of the assay, coordinates of the worm in each field of view were tracked independently. Worm tracks in each field of view were plotted separately using the abovementioned MATLAB software. The end of a worm track in the first field of view was then overlaid with the beginning of the track in the subsequent field of view; this was repeated until all the tracks generated for a given worm formed a single, continuous track.

The images of the iL3 in Figure 1 and the movie of the same iL3 (Movie S1) were generated by recording an EAH435 iL3 on the surface of rat skin using a Zeiss Axio Zoom V16 (Zeiss PlanNeoFluar Z 1x/0.25 FWD 56 mm objective), with the 43 HE dsRed filter, and an attached Basler ace (acA5472-17um) camera until skin penetration was complete. As described above, a series of still images was captured at 4 fps and image acquisition was controlled with the pylon Viewer software (Basler).

### Single-worm skin penetration tracking assays with *S. stercoralis* on human skin

Human skin was obtained either from the forearm of cadaver donors (Accio Biobank Online) or from live donors through plastic surgery. The cadaver donor was a 65-year-old male at death. The cause of death was acute myeloid leukemia. The skin was retrieved from the donor within 8 h after death, frozen, and shipped on dry ice. In the case of live donors, skin was obtained from adult patients (30-50 years old, either male or female) who were undergoing elective plastic surgeries at the UCLA Dermatology Clinic. Fresh skin samples were suspended in 1X PBS immediately after retrieval from the donor. In all cases, the skin, which was normal in appearance, was sectioned (after thawing, in the case of the cadaver donors) and frozen at -80°C. The frozen skin was thawed slowly overnight at 4°C and then allowed to equilibrate to room temperature for 1-2 h before performing skin penetration assays.

On the day of the skin penetration assay, transgenic *S. stercoralis* iL3s were isolated using a Baermann apparatus^43^, as previously described. We then prepared the thawed skin samples. Infective larvae usually come in contact with human hosts when humans are walking barefoot through contaminated soil^9^. To mimic the thinner skin of the dorsum of the foot^62,63^, which appears to be a major region of entry for skin-penetrating nematodes^64-66^, as well as any abrasions that might be caused in this area by walking barefoot, we first exfoliated the human skin samples using an Amope Pedi Perfect electronic foot file that was fitted with the head roller attachment that had ultra coarse grains (#4) for 5-15 s. The exfoliated skin was then placed between cotton pads that were pre-moistened with 1X PBS in a 6 cm Petri plate (Tritech Research, T3315) and used for assays within 1 h of exfoliation. The cotton pad from the top of the skin was removed and the skin surface was lightly blotted with a tissue wipe (VWR International, 82003-820) just prior to the start of the assay. Immediately after blotting, individual worms were placed on the skin surface and time-lapse images were acquired for 10 minutes afterward or until penetration was complete, as described above for the rat skin assays. Up to 4 worms were assayed on the same piece of skin consecutively, and the skin surface was re-moistened with 1X PBS and blotted lightly between each worm. Whenever wild-type and mutant worms were assayed on the same day, assays were performed blind to genotype and blinding was lifted after the experiment was over.

### Single-worm skin penetration tracking assays with *A. ceylanicum* on rat skin

Skin penetration assays with *A. ceylanicum* iL3s were done essentially as described above, with a few differences. *A. ceylanicum* iL3s were typically collected from plates that were 10-14 days old. Worms were stained with DiI, as above, for 10 min and then immediately spun down at 3,000 g for 1 min. All iL3s were plated on 10 cm NGM plates^59^ and allowed to exsheath for 10-15 min. Since DiI strongly stained the sheath, exsheathed worms could be easily identified as the non-fluorescent worms using a Leica M165 FC microscope (ET-mCherry filter set, Leica, 10450195). The exsheathed worms were picked and placed into a watch glass that had 1 mL of BU^58^ pre-mixed with 10 µL of DiI. Approximately 10 min later, individual iL3s were pipetted from the watch glass onto a 10 cm NGM plate^59^ and then transferred onto skin using a paintbrush. Images were captured, as detailed above, for 5 min or until iL3s had penetrated the skin. Only iL3s that were in the second DiI stain for less than an hour were used for skin penetration assays.

### Treatment of worms with haloperidol and dopamine

Haloperidol-treatment of *S. stercoralis*, *S. ratti,* and *A. ceylanicum* iL3s was done overnight at room temperature. Stock solutions of either 20 mM or 40 mM haloperidol (Millipore Sigma, 52-86-8) in DMSO (Millipore Sigma, 67-68-5) were made fresh prior to each experiment. *S. stercoralis* iL3s were treated with 1.5 mM haloperidol in BU^44^, whereas *S. ratti* and *A. ceylanicum* iL3s were treated with 160 µM haloperidol in BU^44^. Vehicle-only controls were treated overnight with an equal concentration of DMSO only; the concentrations of DMSO were 0.8%, 3.8% and 0.4% for the *S. ratti*, *S. stercoralis,* and *A. ceylanicum* iL3s, respectively. Skin penetration assays were then performed as described above.

Treatment of iL3s with dopamine (DA) was done for 1-2 h. A stock solution of 1 M DA (Millipore Sigma, 62-31-7) in ddH_2_O was made fresh on the morning of each assay day and stored in the dark at 4°C until addition to the worm suspension. To expose the worms to DA, the stock solution of DA was added to the worms (that were being treated either with haloperidol or DMSO) to a final concentration of 10 mM. Skin penetration assays were performed as described above.

### Generation of a stable mutant lines using CRISPR/Cas9-mediated mutagenesis

The generation of stable mutant lines was done as previously described^32^ (Fig. S4F). To generate the *Ss-cat-2* mutant line, *S. stercoralis* free-living females were collected from fecal-charcoal plates kept at 25°C for one day or 20°C for two days and then microinjected^67^ with one of the two following injection mixes: mix 1 (for generation of red worms), which had pRP23 (80 ng/µL), pRP31 (80 ng/µL) and pPV540 (50 ng/µL); or mix 2 (for generation of green worms), which had pRP23 (80 ng/µL), pRP44 (80 ng/µL) and pPV540 (50 ng/µL). F_1_ iL3s were isolated from fecal-charcoal plates using a Baermann apparatus^43^ between 5-7 days later. Approximately 100-200 iL3s were plated at a time on 6 cm NGM plates that were seeded with *E. coli* OP50^59^ and then screened for full-body mScarlet-I expression or full-body GFP expression using the Leica M165 FC microscope, with the ET-mCherry filter (Leica, 10450195) or the ET GFP filter set (Leica, 10447408), respectively. The transgenic F_1_ iL3s were picked, pooled and then activated by incubating them, for ∼42 hours, in Dulbecco’s Modified Eagle Medium (DMEM, Gibco, 11995065) at 37°C and 5% CO_2_, as described previously^68^. After incubation, iL3s were collected, washed 3 times in 1X PBS, re-suspended in 200 µL of 1X PBS, and then introduced into a single gerbil by oral gavage. Feces and iL3s were collected from the infected gerbil as described above. The presence of transgenic F_2_/F_3_ iL3s was confirmed by screening, as described above, and dual-colored worms that expressed both mScarlet-I and GFP were picked and used either for skin penetration assays, genotyping, or maintenance of the mutant strain.

A similar approach was used to generate the *Ss-trp-4* mutant line^32^. Free-living females were microinjected^67^ with one of the two mixes: mix 1 (for generation of red worms), which had pRP41 (40 ng/µL), pRP42 (40 ng/µL) pRP40 (80 ng/µL) and pPV540 (50 ng/µL); or mix 2 (for generation of blue worms), which had pRP41 (40 ng/µL), pRP42 (40 ng/µL), pRP50 (80 ng/µL) and pPV540 (50 ng/µL). After 5-7 days, F_1_ iL3s were isolated^43^ and screened either for full-body mScarlet-I signal or full-body Electra2 signal, using the Leica M165 FC microscope, with the ET-mCherry filter (Leica, 10450195) or the ET BFP2 filter set (Leica, 10450571), respectively. Worms were picked, pooled, and introduced into a gerbil by oral gavage as described above.

### Genotyping worms from CRISPR/Cas9 assays

Worm lysis was done as previously described, wherein individual iL3s were placed in 6 µL of worm lysis buffer (50 mM KCl, 10 mM Tris pH 8, 2.5 mM MgCl_2_, 0.45% Nonidet-P40, 0.45% Tween-20, and 0.01% gelatin in ddH_2_O) supplemented with ∼0.12 μg/μL Proteinase-K (Millipore Sigma, 39450-01-6) and ∼1.7% 2-mercaptoethanol (Millipore Sigma, 60-24-2)^55^.

For genotyping at the *Ss-cat-2* locus, each lysed worm was used for 3 PCR reactions: 1) a positive control reaction with primers SG78 and SG80 that target the *Ss-act-2* gene and produce a 416 bp amplicon; 2) a reaction with primers RP32 and RP33, which produce a 666 bp band specifically with the wild-type *Ss-cat-2* allele; and 3) a reaction with primers RP30 and RP34, which produce a 690 bp band specifically with the mutant *Ss-cat-2* allele. For genotyping at the *Ss-trp-4* locus, each lysed worm was similarly used for 3 PCR reactions: 1) the same positive control reaction as above that targets the *Ss-act-2* gene; 2) a reaction with primers RP39 and RP46, which produce a 604 bp band specifically with the wild-type *Ss-trp-4* allele; and 3) a reaction with primers RP39 and RP28, which produce a 720 bp band specifically with the mutant *Ss-trp-4* allele. Genotyping primers are listed in Table S1. The polymerase PlatTaq (Thermo Fisher Scientific, 10966034) was used, and each reaction had a final volume of 25 µL. The PCR reactions were run on an Eppendorf Mastercycler Nexus Gradient (Millipore Sigma, EP6331000025) using the following cycling conditions: initial denaturation 94°C (2 min); 94°C (30 s), 53°C (30 s), 68°C (1 min) x35 cycles; final extension 68°C (5 min); 10°C (hold). PCR products were run out on a 2% agarose gel and stained with GelGreen (Biotium, 41005); the size of each product was gauged by comparing with a 100 bp ladder (New England Biolabs, N3231). The gels were imaged in a ChemiDoc MP Imaging System (Bio-Rad Laboratories) using an exposure time of 1 s. Images were acquired using Image Lab 5.1 (Bio-Rad Laboratories). The presence of bands in each lane was determined using the Lane and Bands tool with the Band Detection Sensitivity set to 100%.

### Histamine assays

*S. stercoralis* free-living females were collected from fecal-charcoal plates kept at 25°C for one day or 20°C for two days and microinjected with pAGR02 at 80 ng/µL using well-established techniques^67^. F_1_ iL3s were isolated from fecal-charcoal plates using a Baermann apparatus^43^ 5-7 days later. Approximately 100-200 iL3s were plated at a time on 6 cm NGM plates that were seeded with *Escherichia coli* OP50^59^ and then screened for mScarlet-I expression in the DA neurons using a Leica M165 FC microscope with the ET-mCherry filter (Leica, 10450195). Only transgenic iL3s with mScarlet-I signal visible in multiple *Ss*-CEP neurons and the *Ss*-ADE neurons at 20-25X magnification were picked for skin penetration assays; expression of the construct in the *Ss*-PDE neurons was variable, so these neurons were likely not always silenced in our experiments. The transgenic F_1_ iL3s were picked and placed in 1 mL of BU saline^58^ and left at 23°C overnight. The following day, transgenic worms were split into two batches: one batch was treated with histamine dihydrochloride (stock concentration = 1 M in ddH_2_O, Millipore Sigma, 56-92-8) diluted to a final concentration of 50 mM in BU^44^; the other batch was treated with an equal volume of ddH_2_O (the solvent for the histamine stock solution), which was also mixed with BU^44^. Skin penetration assays were performed, as described above, 4 h later. Assays were performed blind to experimental condition and blinding was lifted after all the worms had been recorded.

### Fluorescence microscopy

Microscopy of worms was performed using previously established methods for fluorescence microscopy of paralyzed nematodes^44^. *S. stercoralis* free-living females were microinjected, as detailed above, and recovered on fecal-charcoal plates. Transgenic F_1_ iL3s were isolated from these plates after 5-7 days by screening under the Leica M165 FC microscope using either the ET GFP filter set (Leica, 10447408) or the ET mCherry filter set (Leica, 10450195); iL3s were paralyzed with 1% nicotine (Millipore Sigma, 54-11-5) prior to screening. The transgenic iL3s were then exposed to 50 mM levamisole (Millipore Sigma, 16595-80-5) in BU saline^58^, mounted on a slide with 5% Noble agar dissolved in BU^58^, and covered with a coverslip.

Epifluorescence and DIC images were taken with either a 20x objective (Plan-Apochromat 20x/0.8 M27; Zeiss) or a 40x oil objective (Plan-Apochromat 40x/1.4 ∞/0.17 Oil DIC (UV) VIS-IR M27; Zeiss) on an inverted Zeiss AxioObserver microscope equipped with a 38 HE filter set for GFP (BP470/40, FT495, BP 525/50), a 63 HE filter set for mScarlet-I (BP572/25, FT590, BP629/62), a 96 HE filter set for Electra2 (BP 390/40, FT420, BP450/40), and a Hamamatsu ORCA-Flash 4.0 camera; fluorescence illumination was provided by Colibri 7 LEDs (LED-Module 475 nm). All images were captured using Zeiss ZEN 2 (blue edition) software. Images in magenta were pseudo-colored in Fiji 2.9.0/1.53t^60^ and image montages were generated in Adobe Photoshop 25.5.0 and Adobe Illustrator 28.4.1.

### Statistical analysis

Statistical analyses were performed in Prism 10.0.0. The statistical tests used for each experiment are listed in the figure legends; two-tailed tests were used for all statistical analyses. Non-parametric tests were used when the data were found to be non-parametrically distributed, as determined by tests for normality in Prism. Sample sizes were determined by power analysis using G*Power 3.1.9.6. All statistical analyses are listed in Dataset S1.

## ACKNOWLEDGMENTS

We thank Navonil Banerjee, Michelle Castelletto, and Breanna Walsh for thoughtful comments on the manuscript and Tiffany Mao for hand-drawn illustrations. The illustrations shown in Fig. 1 and Fig. S1, S2, S4F, and S6A were created with BioRender. The gene models of *Ce-trp-4* and *Ss-trp-4* in Fig. 7D were adapted from models made using the Exon-Intron Graphic Maker (http://www.wormweb.org/exonintron). This work was supported by NIH F32AI174816 (R.P.), NIH MARC T34GM008563 (A.G.R.), funds provided by the University of Washington School of Medicine and NIH DP2AI184544 (A.S.B.), NIH R01AR081337 (G.W.A), and NIH R01AI175183 (E.A.H.).

## REFERENCES

1 Riaz, M. et al. Prevalence, risk factors, challenges, and the currently available diagnostic tools for the determination of helminths infections in human. EJI 18, 1–15, doi:10.1177/2058739220959915 (2020).

2 Buonfrate, D. et al. The global prevalence of *Strongyloides stercoralis* infection. Pathogens 9, doi:10.3390/pathogens9060468 (2020).

3 WHO. Soil-transmitted helminth infections. (2023).

4 McClure, C. R., Patel, R. & Hallem, E. A. Invade or die: behaviours and biochemical mechanisms that drive skin penetration in *Strongyloides* and other skin-penetrating nematodes. Philos Trans R Soc Lond B Biol Sci 379, 20220434, doi:10.1098/rstb.2022.0434 (2024).

5 Forrer, A., et al. *Strongyloides stercoralis* is associated with significant morbidity in rural Cambodia, including stunting in children. PLoS Negl Trop Dis 11, e0005685, doi:10.1371/journal.pntd.0005685 (2017).

6 Fauziah, N., Ar-Rizqi, M. A., Hana, S., Patahuddin, N. M. & Diptyanusa, A. Stunting as a risk factor of soil-transmitted helminthiasis in children: a literature review. Interdiscip Perspect Infect Dis 2022, 8929025, doi:10.1155/2022/8929025 (2022).

7 Shang, Y. et al. Stunting and soil-transmitted-helminth infections among school-age pupils in rural areas of southern China. Parasit Vectors 3, 97, doi:10.1186/1756-3305-3-97 (2010).

8 Czeresnia, J. M. & Weiss, L. M. Strongyloides stercoralis. Lung 200, 141–148, doi:10.1007/s00408-022-00528-z (2022).

9 Gordon, C. A. et al. Strongyloidiasis. Nat Rev Dis Primers 10, 6, doi:10.1038/s41572-023-00490-x (2024).

10 Maroto, R. et al. First report of anthelmintic resistance in gastrointestinal nematodes of sheep from Costa Rica. Vet Med Int 2011, 145312, doi:10.4061/2011/145312 (2011).

11 Zaman, V., Dawkins, H. J. & Grove, D. I. Scanning electron microscopy of the penetration of newborn mouse skin by *Strongyloides ratti* and *Ancylostoma caninum* larvae. Southeast Asian J Trop Med Public Health 11, 212–219 (1980).

12 Viney, M. & Kikuchi, T. *Strongyloides ratti* and *S. venezuelensis* - rodent models of *Strongyloides* infection. Parasitology 144, 285–294, doi:10.1017/S0031182016000020 (2017).

13 Viney, M. E. & Lok, J. B. The biology of *Strongyloides* spp. WormBook, 1–17, doi:10.1895/wormbook.1.141.2 (2015).

14 Buonfrate, D., Bradbury, R. S., Watts, M. R. & Bisoffi, Z. Human strongyloidiasis: complexities and pathways forward. Clin Microbiol Rev 36, e0003323, doi:10.1128/cmr.00033-23 (2023).

15 Patel, R. et al. The generation of stable transgenic lines in the human-infective nematode *Strongyloides stercoralis*. G3 (Bethesda) 14, doi:10.1093/g3journal/jkae122 (2024).

16 Schultz, R. D. & Gumienny, T. L. Visualization of *Caenorhabditis elegans* cuticular structures using the lipophilic vital dye DiI. J Vis Exp, e3362, doi:10.3791/3362 (2012).

17 Matoltsy, A. G. in Biology of the Integument 2: Vertebrates Ch. 15, 272–277 (1984).

18 Sawin, E. R., Ranganathan, R. & Horvitz, H. R. *C. elegans* locomotory rate is modulated by the environment through a dopaminergic pathway and by experience through a serotonergic pathway. Neuron 26, 619–631, doi:10.1016/s0896-6273(00)81199-x (2000).

19 Han, B. et al. Dopamine signaling tunes spatial pattern selectivity in *C. elegans*. eLife 6, doi:10.7554/eLife.22896 (2017).

20 Fok, A. et al. High-fidelity encoding of mechanostimuli by tactile food-sensing neurons requires an ensemble of ion channels. Cell Rep 42, 112452, doi:10.1016/j.celrep.2023.112452 (2023).

21 Tanimoto, Y., et al. *In actio* optophysiological analyses reveal functional diversification of dopaminergic neurons in the nematode *C. elegans*. Sci Rep 6, 26297, doi:10.1038/srep26297 (2016).

22 Krum, B. N. et al. Haloperidol interactions with the DOP-3 receptor in *Caenorhabditis elegans*. Mol Neurobiol 58, 304–316, doi:10.1007/s12035-020-02124-9 (2021).

23 Sanyal, S. et al. Dopamine modulates the plasticity of mechanosensory responses in *Caenorhabditis elegans*. EMBO J 23, 473–482, doi:10.1038/sj.emboj.7600057 (2004).

24 Lints, R. & Emmons, S. W. Patterning of dopaminergic neurotransmitter identity among *Caenorhabditis elegans* ray sensory neurons by a TGFbeta family signaling pathway and a Hox gene. Development 126, 5819–5831, doi:10.1242/dev.126.24.5819 (1999).

25 Molinoff, P. B. & Axelrod, J. Biochemistry of catecholamines. Annu Rev Biochem 40, 465–500, doi:10.1146/annurev.bi.40.070171.002341 (1971).

26 Jayanthi, L. D. et al. The *Caenorhabditis elegans* gene T23G5.5 encodes an antidepressant- and cocaine-sensitive dopamine transporter. Mol Pharmacol 54, 601–609 (1998).

27 Wang, J. et al. The conserved domain database in 2023. Nucleic Acids Res 51, D384–D388, doi:10.1093/nar/gkac1096 (2023).

28 Altun, Z. F. & Hall, D. H. Nervous system, neuronal support cells. WormAtlas, doi:doi:10.3908/wormatlas.1.19 (2012).

29 Sulston, J., Dew, M. & Brenner, S. Dopaminergic neurons in the nematode *Caenorhabditis elegans*. J Comp Neurol 163, 215–226, doi:10.1002/cne.901630207 (1975).

30 Bryant, A. S., DeMarco, S. F. & Hallem, E. A. *Strongyloides* RNA-seq Browser: a web-based software platform for on-demand bioinformatics analyses of *Strongyloides* species. G3 (Bethesda) 11, doi:10.1093/g3journal/jkab104 (2021).

31 Hunt, V. L. et al. The genomic basis of parasitism in the *Strongyloides* clade of nematodes. Nat Genet 48, 299–307, doi:10.1038/ng.3495 (2016).

32 Banerjee, N. et al. Carbon dioxide shapes parasite-host interactions in a human-infective nematode. Curr Biol (2024).

33 Pokala, N., Liu, Q., Gordus, A. & Bargmann, C. I. Inducible and titratable silencing of *Caenorhabditis elegans* neurons *in vivo* with histamine-gated chloride channels. Proc Natl Acad Sci U S A 111, 2770–2775, doi:10.1073/pnas.1400615111 (2014).

34 Kang, L., Gao, J., Schafer, W. R., Xie, Z. & Xu, X. Z. *C. elegans* TRP family protein TRP-4 is a pore-forming subunit of a native mechanotransduction channel. Neuron 67, 381–391, doi:10.1016/j.neuron.2010.06.032 (2010).

35 Zhang, M. et al. TRP (transient receptor potential) ion channel family: structures, biological functions and therapeutic interventions for diseases. Signal Transduct Target Ther 8, 261, doi:10.1038/s41392-023-01464-x (2023).

36 Blaxter, M. L. et al. A molecular evolutionary framework for the phylum Nematoda. Nature 392, 71–75, doi:10.1038/32160 (1998).

37 Ashton, F. T., Bhopale, V. M., Fine, A. E. & Schad, G. A. Sensory neuroanatomy of a skin-penetrating nematode parasite: *Strongyloides stercoralis*. I. Amphidial neurons. J Comp Neurol 357, 281–295, doi:10.1002/cne.903570208 (1995).

38 Fine, A. E., Ashton, F. T., Bhopale, V. M. & Schad, G. A. Sensory neuroanatomy of a skin-penetrating nematode parasite *Strongyloides stercoralis*. II. Labial and cephalic neurons. J Comp Neurol 389, 212–223 (1997).

39 Chase, D. L., Pepper, J. S. & Koelle, M. R. Mechanism of extrasynaptic dopamine signaling in *Caenorhabditis elegans*. Nat Neurosci 7, 1096–1103, doi:10.1038/nn1316 (2004).

40 Kindt, K. S. et al. Dopamine mediates context-dependent modulation of sensory plasticity in *C. elegans*. Neuron 55, 662–676, doi:10.1016/j.neuron.2007.07.023 (2007).

41 Ezak, M. J. & Ferkey, D. M. The *C. elegans* D2-like dopamine receptor DOP-3 decreases behavioral sensitivity to the olfactory stimulus 1-octanol. PLoS One 5, e9487, doi:10.1371/journal.pone.0009487 (2010).

42 Castelletto, M. L. et al. Diverse host-seeking behaviors of skin-penetrating nematodes. PLoS Pathog 10, e1004305, doi:10.1371/journal.ppat.1004305 (2014).

43 Lok, J. B. *Strongyloides stercoralis*: a model for translational research on parasitic nematode biology. WormBook, 1-18, doi:10.1895/wormbook.1.134.1 (2007).

44 Bryant, A. S., Ruiz, F., Lee, J. H. & Hallem, E. A. The neural basis of heat seeking in a human-infective parasitic worm. Curr Biol 32, 2206–2221 e2206, doi:10.1016/j.cub.2022.04.010 (2022).

45 Bryant, A. S., Akimori, D., Stoltzfus, J. D. C. & Hallem, E. A. A standard workflow for community-driven manual curation of *Strongyloides* genome annotations. Philos Trans R Soc Lond B Biol Sci 379, 20220443, doi:10.1098/rstb.2022.0443 (2024).

46 Nguyen, L. T., Schmidt, H. A., von Haeseler, A. & Minh, B. Q. IQ-TREE: a fast and effective stochastic algorithm for estimating maximum-likelihood phylogenies. Mol Biol Evol 32, 268–274, doi:10.1093/molbev/msu300 (2015).

47 Hoang, D. T., Chernomor, O., von Haeseler, A., Minh, B. Q. & Vinh, L. S. UFBoot2: Improving the Ultrafast Bootstrap Approximation. Mol Biol Evol 35, 518–522, doi:10.1093/molbev/msx281 (2018).

48 Letunic, I. & Bork, P. Interactive Tree Of Life (iTOL): an online tool for phylogenetic tree display and annotation. Bioinformatics 23, 127–128, doi:10.1093/bioinformatics/btl529 (2007).

49 Marchler-Bauer, A. & Bryant, S. H. CD-Search: protein domain annotations on the fly. Nucleic Acids Res 32, W327–331, doi:10.1093/nar/gkh454 (2004).

50 Stoltzfus, J. D., Minot, S., Berriman, M., Nolan, T. J. & Lok, J. B. RNAseq analysis of the parasitic nematode *Strongyloides stercoralis* reveals divergent regulation of canonical dauer pathways. PLoS Negl Trop Dis 6, e1854, doi:10.1371/journal.pntd.0001854 (2012).

51 Xiao, R. & Xu, X. Z. *C. elegans* TRP channels. Adv Exp Med Biol 704, 323–339, doi:10.1007/978-94-007-0265-3_18 (2011).

52 Kahn-Kirby, A. H. & Bargmann, C. I. TRP channels in *C. elegans*. Annu Rev Physiol 68, 719–736, doi:10.1146/annurev.physiol.68.040204.100715 (2006).

53 Goodman, M. B. Mechanosensation. WormBook, 1-14, doi:10.1895/wormbook.1.62.1 (2006).

54 Venkatachalam, K. & Montell, C. TRP channels. Annu Rev Biochem 76, 387–417, doi:10.1146/annurev.biochem.75.103004.142819 (2007).

55 Gang, S. S. et al. Targeted mutagenesis in a human-parasitic nematode. PLoS Pathog 13, e1006675, doi:10.1371/journal.ppat.1006675 (2017).

56 Doench, J. G. et al. Optimized sgRNA design to maximize activity and minimize off-target effects of CRISPR-Cas9. Nat Biotechnol 34, 184–191, doi:10.1038/nbt.3437 (2016).

57 Papadaki, S. et al. Dual-expression system for blue fluorescent protein optimization. Sci Rep 12, 10190, doi:10.1038/s41598-022-13214-0 (2022).

58 Hawdon, J. M. & Schad, G. A. Long-term storage of hookworm infective larvae in buffered saline solution maintains larval responsiveness to host signals. J Helm Soc Wash 58, 140–142 (1991).

59 Stiernagle, T. Maintenance of *C. elegans*. WormBook, 1–11, doi:10.1895/wormbook.1.101.1 (2006).

60 Schindelin, J. et al. Fiji: an open-source platform for biological-image analysis. Nat Methods 9, 676–682, doi:10.1038/nmeth.2019 (2012).

61 Tinevez, J. Y. et al. TrackMate: an open and extensible platform for single-particle tracking. Methods 115, 80–90, doi:10.1016/j.ymeth.2016.09.016 (2017).

62 Lee, Y. & Hwang, K. Skin thickness of Korean adults. Surg Radiol Anat 24, 183–189, doi:10.1007/s00276-002-0034-5 (2002).

63 Oltulu, P., Ince, B., Kokbudak, N., Findik, S. & Kilinc, F. Measurement of epidermis, dermis, and total skin thicknesses from six different body regions with a new ethical histometric technique. Turk J Plast Surg 26, 56–61, doi:10.4103/tjps.TJPS_2_17.

64 Hall, A. D., Luckett, K. M. & Williams, K. M. Bullous cutaneous larva migrans of the foot. Am J Trop Med Hyg 110, 625–626, doi:10.4269/ajtmh.23-0750 (2024).

65 Green, R., Somayaji, R. & Chia, J. C. Bullous cutaneous larva migrans. CMAJ 195, E1040, doi:10.1503/cmaj.230583 (2023).

66 Gomez-Moyano, E., Pilar, L. M., Simonsen, S. B. & Vera-Casano, A. A serpiginous, itchy rash on the foot. Cleve Clin J Med 83, 494–495, doi:10.3949/ccjm.83a.15110 (2016).

67 Castelletto, M. L. & Hallem, E. A. Generating transgenics and knockouts in *Strongyloides* species by microinjection. J Vis Exp, doi:10.3791/63023 (2021).

68 Gang, S. S. et al. Chemosensory mechanisms of host seeking and infectivity in skin-penetrating nematodes. Proc Natl Acad Sci U S A 117, 17913–17923, doi:10.1073/pnas.1909710117 (2020).

