## Supplemental Material for "Dopamine signaling drives skin invasion by human-infective nematodes"

**SUPPLEMENTAL FIGURES**

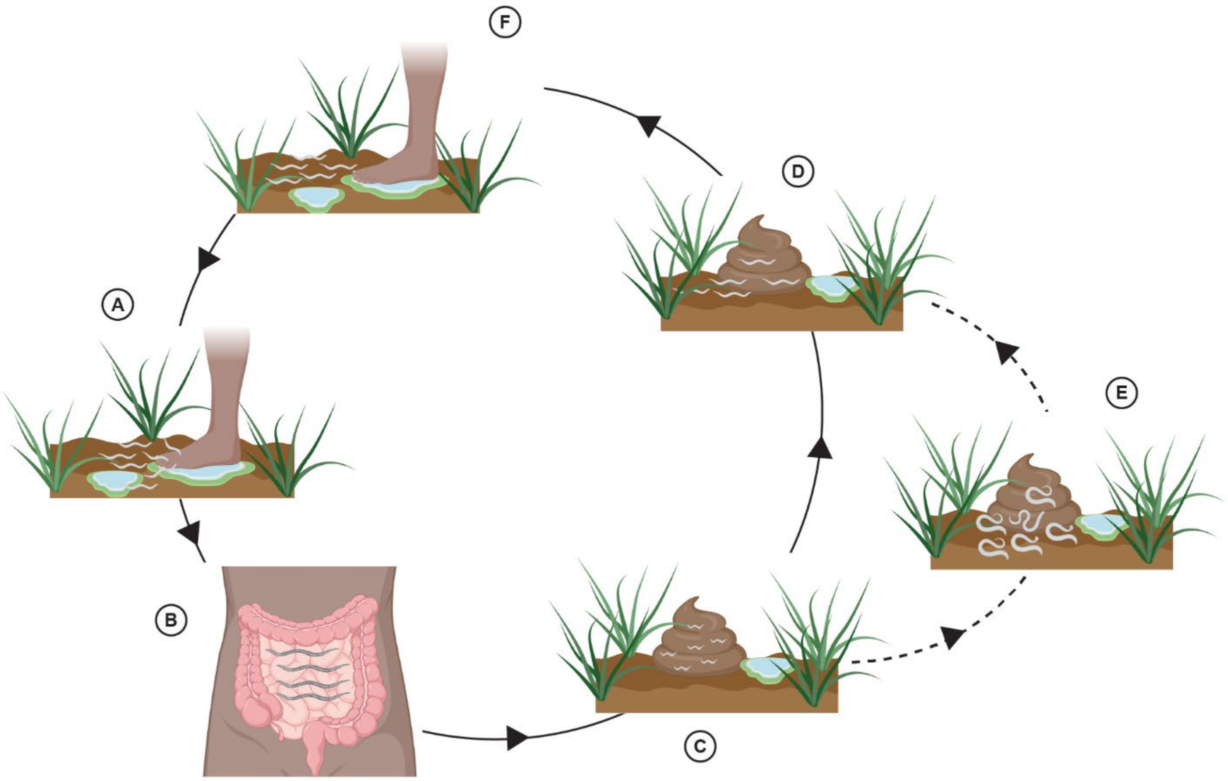


**Figure S1. The life cycle of *S. stercoralis.*** The life cycle of *S. stercoralis* is complex and composed of both intra-host and extra-host life stages[1]. The intra-host portion of the life cycle begins when infective third-stage larvae (iL3s) penetrate the skin of a host and enter the body (A); the natural hosts of *S. stercoralis* include humans (as depicted in the figure), some non-human primates, and dogs[2,3]. Development is paused in iL3s and resumes upon host entry[4]. Following skin penetration, larvae travel through the body and reach the host duodenal mucosa, where they live and reproduce as parasitic adults[5] (B). Parasitic adults lay eggs in the duodenal mucosa[5], which hatch into post-parasitic, first-stage (L1) larvae. A subset of the post-parasitic L1s develop into autoinfective third-stage larvae (aL3s), which can reinfect the host and perpetuate the *S. stercoralis* infection (not shown). Alternatively, post-parasitic L1s are released from the host, into the surrounding environment, in feces (C). Outside of the host, post-parasitic L1s are fated to one of two developmental routes: they either develop directly into iL3s (D) or they develop into free-living females and males (E). All the progeny of free-living females and males become iL3s. The iL3s actively seek out new hosts, using cues such as heat and host-emitted odorants[2,6] (F). The life cycle of *S. ratti* is very similar to *S. stercoralis*, except that *S. ratti* infects rats and all the progeny of parasitic adults exit the host as eggs (thus, *S. ratti* does not have an autoinfective cycle).

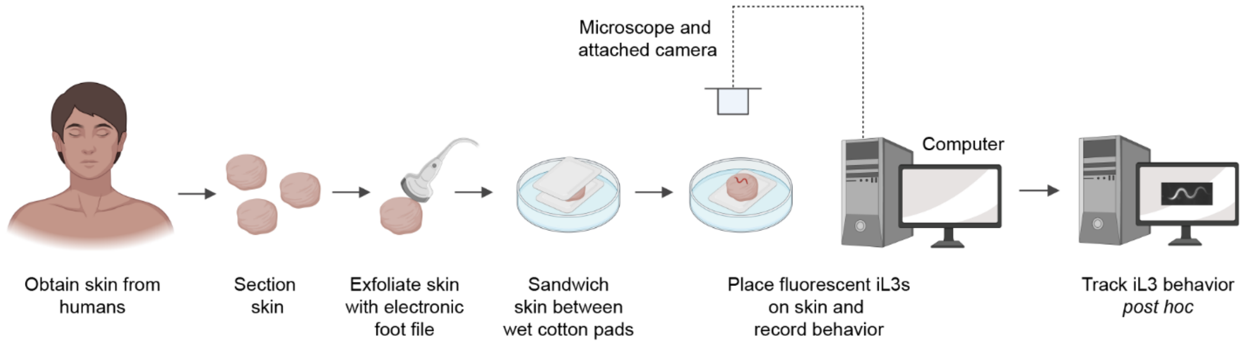


**Figure S2. An *ex vivo* assay for studying skin-penetration behaviors on human skin.** Skin samples are obtained either from cadaver donors (forearm skin) or from patients that underwent surgery (breast or abdominal skin) and then sectioned. Humans often come in contact with *S. stercoralis* when walking barefoot through contaminated soil[7]. The iL3s then penetrate, often through skin on the top surface of the foot or the toes[8-10], and enter the body. To mimic the relative thinness of the skin from the top surface of the foot and toes[11,12], as well as any micro-abrasions caused by walking barefoot in soil, we file the human skin in our assays for 5-15 s with an electronic foot file. We then sandwich the sectioned, filed skin between cotton pads that were pre-moistened with 1X PBS to maintain moisture. We place fluorescent iL3s on the skin surface and acquire time-lapse images of behavior for 10 min thereafter or until penetration is complete. Time-lapse images are recorded using a fluorescence microscope and camera, and videos are analyzed *post hoc*.

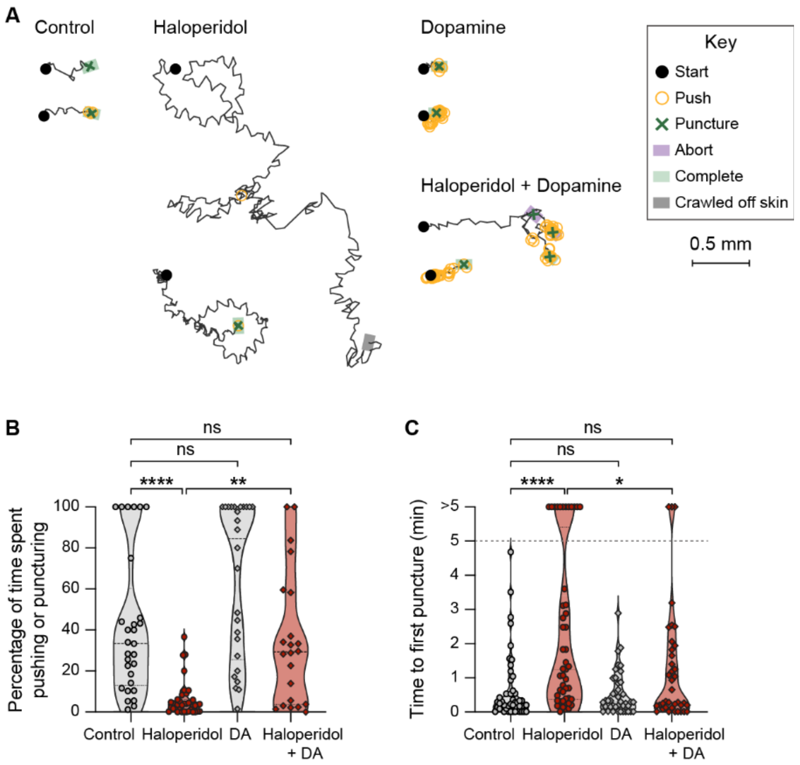


**Figure S3. Pharmacological manipulation of dopamine signaling blocks skin penetration in *S. ratti*. A.** Haloperidol inhibits skin penetration and dopamine (DA) rescues this phenotype in *S. ratti* iL3s. Tracks show two representative worms from each group; the haloperidol-treated group shows one representative worm that completed penetration and one that neither punctured nor completed penetration. The haloperidol-treated worm that did not complete penetration crawled off the edge of the skin into the underlying saline, as depicted by a gray box. Representative worms were chosen as detailed in Fig. 3A. The key shows the behavioral motifs that were tracked. **B.** Haloperidol reduces the percentage of time that *S. ratti* iL3s spend pushing and puncturing the skin, and this effect is rescued by addition of exogenous DA. Violin plot depicts the percentage of time that worms from each treatment group spent engaging in pushes or punctures. n = 22-36 iL3s per condition. ****$p<0.0001$, **$p<0.01$, ns = not significant, Kruskal-Wallis test with Dunn's post-test. The iL3s that had initiated penetration by the time the recording started were excluded from this analysis. **C.** Haloperidol delays the time to first puncture and DA rescues this behavioral phenotype. Violin plot depicts the time taken by worms from each treatment group to puncture the skin for the first time since placement on skin. n = 38-48 iL3s per condition. ****$p<0.0001$, *$p<0.05$, ns = not significant, Kruskal-Wallis test with Dunn's post-test. The dotted line at y = 5 indicates the time at which the assay ended; the dots above this line indicate worms that failed to puncture the skin by the end of the assay. For B-C, dots depict individual worms, dashed lines indicate medians, and dotted lines indicate interquartile ranges. Behavioral parameters plotted in B were obtained from 4 independent replicate experiments and those in C were obtained from 5 independent replicate experiments.

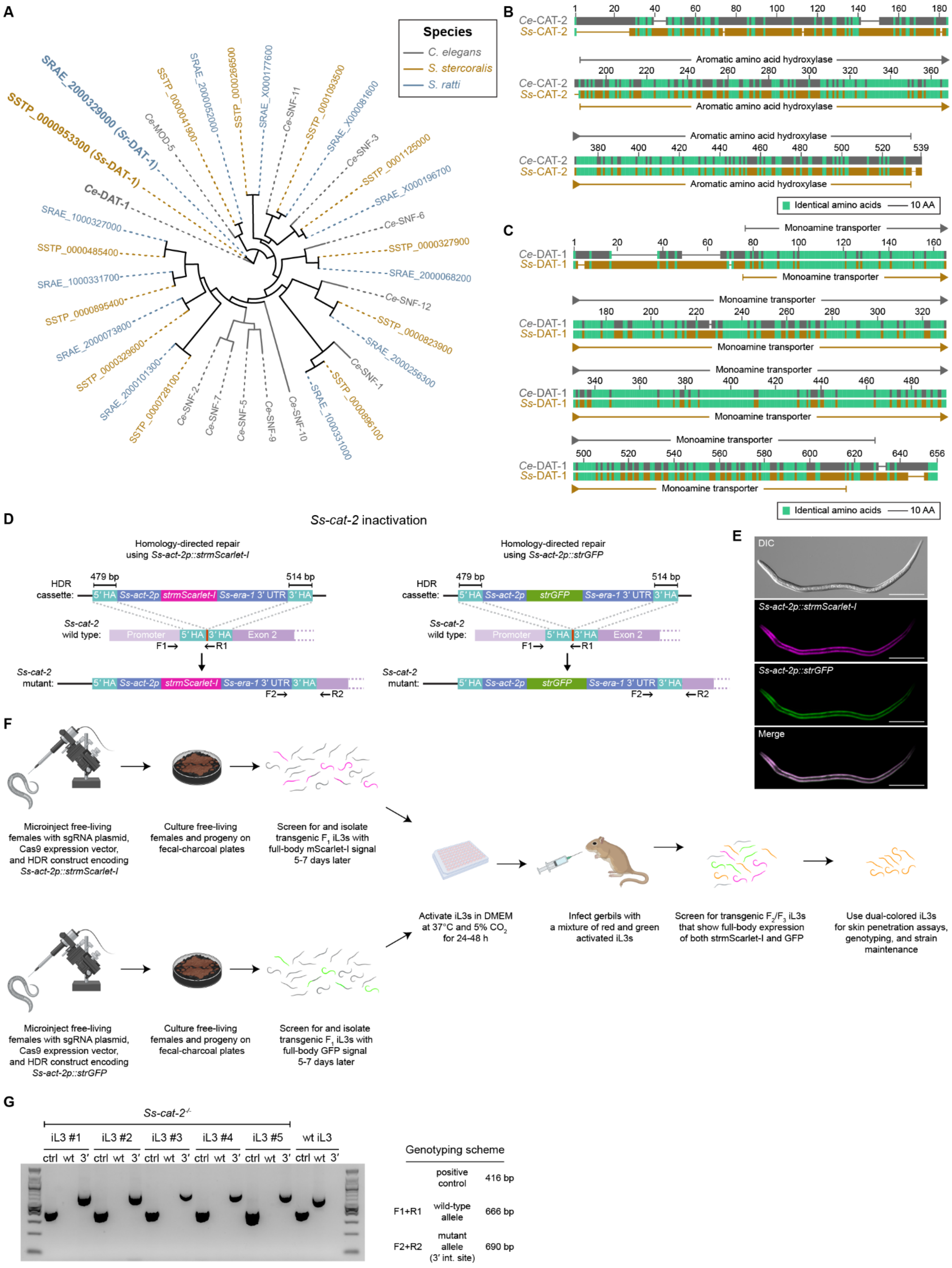

A
Species
C. elegans
S. stercoralis
S. ratti
SRAE_2000329000 (Sr-DAT-1)
SSTP_0000953300 (Ss-DAT-1)
Ce-DAT-1
Ce-MOD-5
Ce-SNF-11
Ce-SNF-3
Ce-SNF-6
Ce-SNF-12
Ce-SNF-1
Ce-SNF-10
Ce-SNF-9
Ce-SNF-5
Ce-SNF-7
Ce-SNF-2
B
Ce-CAT-2
Ss-CAT-2
Aromatic amino acid hydroxylase
Identical amino acids
10 AA
C
Ce-DAT-1
Ss-DAT-1
Monoamine transporter
Identical amino acids
10 AA
D
Ss-cat-2 inactivation
Homology-directed repair
using Ss-act-2p::strmScarlet-I
Homology-directed repair
using Ss-act-2p::strGFP
HDR
cassette:
479 bp
514 bp
5′ HA
Ss-act-2p
strmScarlet-I
strGFP
Ss-era-1 3′ UTR
3′ HA
Ss-cat-2
wild type:
Promoter
Exon 2
F1→
←R1
Ss-cat-2
mutant:
F2→
←R2
E
DIC
Ss-act-2p::strmScarlet-I
Ss-act-2p::strGFP
Merge
F
Microinject free-living females with sgRNA plasmid, Cas9 expression vector, and HDR construct encoding Ss-act-2p::strmScarlet-I
Culture free-living females and progeny on fecal-charcoal plates
Screen for and isolate transgenic F1 iL3s with full-body mScarlet-I signal 5-7 days later
Microinject free-living females with sgRNA plasmid, Cas9 expression vector, and HDR construct encoding Ss-act-2p::strGFP
Culture free-living females and progeny on fecal-charcoal plates
Screen for and isolate transgenic F1 iL3s with full-body GFP signal 5-7 days later
Activate iL3s in DMEM at 37°C and 5% CO2 for 24-48 h
Infect gerbils with a mixture of red and green activated iL3s
Screen for transgenic F2/F3 iL3s that show full-body expression of both strmScarlet-I and GFP
Use dual-colored iL3s for skin penetration assays, genotyping, and strain maintenance
G
Ss-cat-2-/-
iL3 #1
iL3 #2
iL3 #3
iL3 #4
iL3 #5
wt iL3
ctrl wt 3′
Genotyping scheme
positive control 416 bp
F1+R1 wild-type allele 666 bp
F2+R2 mutant allele (3′ int. site) 690 bp

**Figure S4. Inactivation of dopamine signaling in *S. stercoralis* using CRISPR/Cas9-mediated mutagenesis. A.** Phylogenetic analysis shows the closest homologs of *C. elegans* DAT-1 (gray), *S. stercoralis* (brown), and *S. ratti* (blue). Putative homologs in each genome were identified by performing TBLASTN searches of the *C. elegans* DAT-1 protein sequence against either the *S. stercoralis* genome or the *S. ratti* genome in WBPS18. The tree has all known members of the sodium neurotransmitter symporter family (SNF) of proteins in *C. elegans*, of which DAT-1 is a member, and predicted homologs in *S. stercoralis* and *S. ratti*. **B.** Schematic representation of the alignment of the amino acid sequences of the *C. elegans* CAT-2 protein (isoform A, top) and the *S. stercoralis* CAT-2 protein (bottom). The aromatic amino acid hydroxylase domains, which are the catalytic domains of this family of enzymes[13], of the two proteins are 56% identical. **C.** Schematic representation of the alignment of the amino acid sequences of the *C. elegans* DAT-1 protein (top) and the *S. stercoralis* DAT-1 protein (bottom). The monoamine transporter domains are ~69% identical. In **B**-**C**, identical amino acids are shown in green and the domains were identified by searches in the Conserved Domain Database. Drawings are to scale and the scale bar = 10 amino acids (AA). **D.** Inactivation of *Ss-cat-2* was achieved by integration of a transcriptional reporter for the *Ss-act-2* gene at the cut site generated by Cas9. The *Ss-act-2* transcriptional reporter consisted of the promoter of the *Ss-act-2* gene fused with a *Strongyloides*-codon-optimized gene encoding either mScarlet-I (left schematic) or GFP (right schematic) and the *Ss-era-1* 3′ UTR. This transgene was flanked by 5′ and 3′ homology arms (HAs); the 5′ HA matched the 479 bp fragment immediately upstream of the CRISPR site in *Ss-cat-2* (depicted in red) and the 3′ HA matched a 514 bp fragment immediately downstream of this same site. Homology-directed repair following double-stranded breaks generated by Cas9 resulted in insertion of either the *Ss-act-2p::strmScarlet-I::Ss-era-1* 3′ UTR transgene or the *Ss-act-2p::strGFP::Ss-era-1* 3′ UTR transgene into the *Ss-cat-2* locus, creating a stop codon early in the second exon and thereby preventing expression of the mutant allele. The approximate binding sites of the genotyping primers are also shown. The presence of a PCR amplicon from F1 and R1 indicated a wild-type locus, as R1 overlaps the entire Cas9 cut site. The presence of a PCR amplicon from F2 and R2 indicated a mutant locus, as F2 lies in the *Ss-era-1* 3′ UTR and R2 lies in the region of the *Ss-cat-2* gene that is downstream of the 3′ HA. **E.** Expression of the *Ss-act-2p::strmScarlet-I* transgene (magenta) and *Ss-act-2p::strGFP* transgene (green) over the entire body wall muscle of a dual-colored *Ss-cat-2*$^{-/-}$ mutant. The worm is oriented with the ventral side facing downwards and the head to the left. Scale bar = 100 µm. **F**. Schematic shows the approach used to generate a mutant *Ss-cat-2*$^{-/-}$ stable line[14]. *S. stercoralis* free-living females were microinjected with one of the following mixtures: mixture 1, which consists of the Cas9 expression vector, the single guide RNA (sgRNA) expression vector, and a plasmid that provides the template for insertion of *Ss-act-2p::strmScarlet-I* into the *Ss-cat-2* locus via homology-directed repair (HDR) (top left); or mixture 2, which consists of all the components in mixture 1, except that the HDR template had the *Ss-act-2p::strGFP* transgene (bottom left). Between 5-7 days later, transgenic $F_1$ iL3s that expressed either mScarlet-I or GFP across the entire body wall muscle were selected by fluorescence microscopy; iL3s with full-body expression of either fluorescent protein are more likely to have integration of the transgenes into the *Ss-cat-2* locus and stable transmission of the mutant allele to progeny[14,15]. These iL3s were activated by incubating them in a mixture of DMEM and antibiotics at 37°C and 5% $CO_2$ for 24-48 h[16-19]. The activated, transgenic iL3s were then gavaged into a single gerbil orally[14,20]. Three weeks later, transgenic $F_2$/$F_3$ iL3s that expressed both mScarlet-I and GFP were selected by fluorescence microscopy. These dual-colored iL3s had one copy of the *Ss-cat-2* gene replaced with *Ss-act-2p::strmScarlet-I* and the other copy replaced with *Ss-act-2p::strGFP*[58]; thus, they were homozygous *Ss-cat-2* mutants. These homozygous mutants were then used for skin penetration assays and propagation of the strain. **G.** Image shows a representative agarose gel that was loaded and run with amplicons from genotyping PCRs of five *Ss-cat-2*$^{-/-}$ dual-colored iL3s and one wild-type iL3. For each worm, products from the following PCR reactions were loaded in order: a positive control for the PCR reaction (exon 1 of the *Ss-act-2* gene), which produced a 416 bp band (ctrl); a PCR reaction for genotyping the wild-type allele of *Ss-cat-2* (using primers F1 and R1), which produced a 666 bp band (wt) in wild-type iL3s; and a PCR reaction for genotyping the 3′ integration site of the HDR template (using primers F2 and R2), which produced a 690 bp band (3′) in carriers of the mutant *Ss-cat-2* allele. The first and last lanes carried a New England Biolabs 100 bp ladder. “3′ int. site” = 3′ integration site.

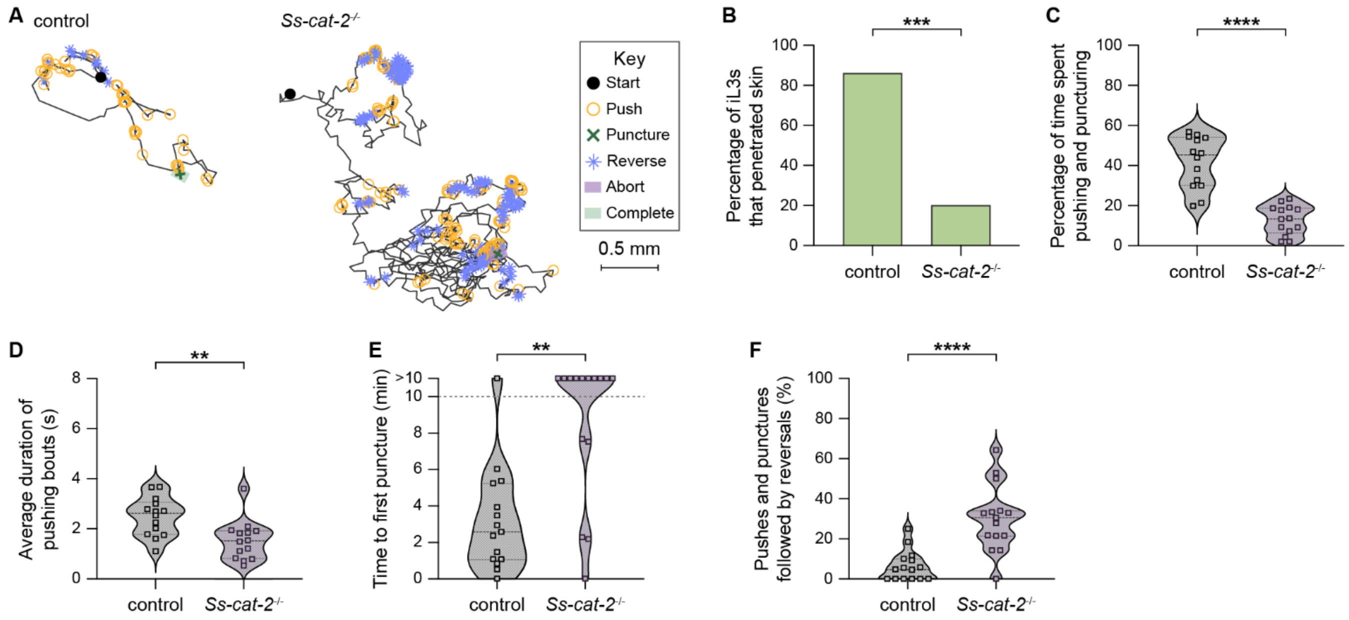


**Figure S5. Inactivation of *Ss-cat-2* impairs penetration of human skin. A.** *Ss-cat-2*$^{-/-}$ iL3s exhibit reduced skin penetration and altered behaviors on human skin. Tracks show the behaviors of representative wild-type iL3 that punctured and completed penetration and a representative *Ss-cat-2*$^{-/-}$ iL3 that punctured but did not complete penetration on human skin. Representative worms were defined as in Fig. 3A. The key details the behavioral motifs that were tracked. **B.** Inactivation of *Ss-cat-2* severely impairs skin penetration. Bar graph shows the percentage of wild-type and *Ss-cat-2*$^{-/-}$ iL3s that completed skin penetration. n = 15 iL3s per genotype. ***$p$<0.001, Fisher's exact test. **C.** *Ss-cat-2*$^{-/-}$ iL3s pushed and punctured the skin for less time than control worms. Violin plot depicts the percentage of time on skin that control and *Ss-cat-2*$^{-/-}$ iL3s spent engaging in pushes or punctures. n = 14 iL3s per genotype. ****$p$<0.0001, unpaired t-test. The iL3s that had initiated penetration by the time the recording started were excluded from this analysis. **D.** The pushing bouts of *Ss-cat-2*$^{-/-}$ iL3s were shorter than those of control iL3s. For each worm, the duration of each individual pushing bout was averaged and then plotted. n = 14 iL3s per genotype. **$p$<0.01, unpaired t-test. **E.** Inactivation of *Ss-cat-2* inhibits punctures. Violin plot depicts the time taken by control vs. *Ss-cat-2*$^{-/-}$ worms to puncture the skin for the first time since placement upon skin. n = 15 iL3s per genotype. **$p$<0.01, Mann-Whitney test. The dotted line at y = 10 indicates the time at which the assay ended; the dots above this line indicate animals that failed to puncture the skin by the end of the assay period. **F.** *Ss-cat-2*$^{-/-}$ iL3s frequently reverse after a push or puncture. Violin plot shows the percentage of pushes or punctures that were followed by backward locomotion that lasted at least 1 s for each genotype. n = 15 iL3s per genotype. ****$p$<0.0001, Mann-Whitney test. For **C**-**F**, dots depict individual worms, dashed lines indicate medians, and dotted lines indicate interquartile ranges. Behavioral parameters plotted in **B**-**F** were obtained from 3 independent replicate experiments, each using skin from a distinct human donor.

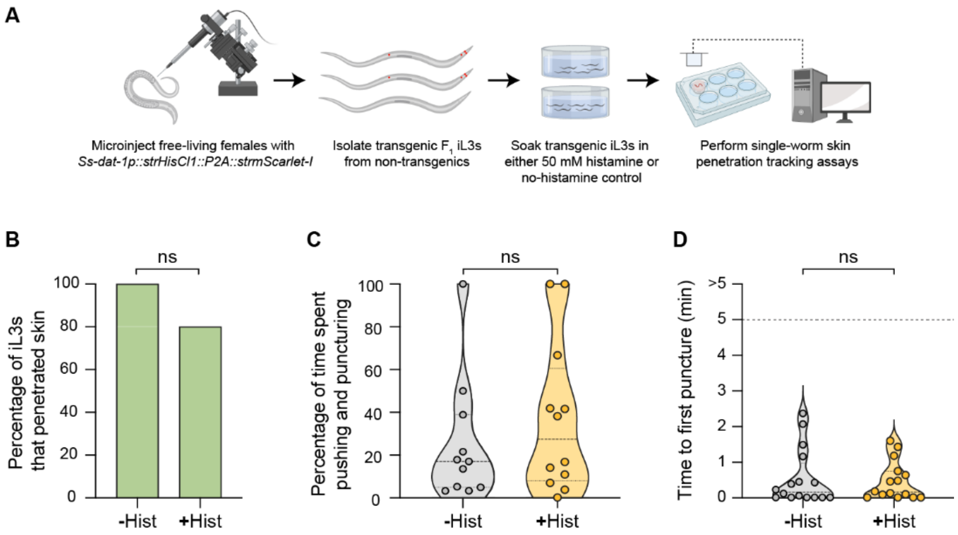


**Figure S6. Chemogenetic silencing of the dopaminergic neurons using the histamine-gated chloride channel HisCl1. A.** Schematic of the approach for silencing the dopaminergic neurons of *S. stercoralis* using HisCl1[21,22]. The gonads of free-living adult females were microinjected with an *Ss-dat-1p::strHisCl1::P2A::strmScarlet-I* transgene using standard techniques[23]. Transgenic $F_1$ iL3s that expressed both HisCl1 and mScarlet-I in the dopaminergic neurons were isolated by performing fluorescence microscopy-based screening for mScarlet-I signal. The transgenics were then separated into two groups: one group was treated with 50 mM histamine and the other group was treated with the vehicle only ($ddH_2O$). Skin penetration assays were then performed as detailed in Fig. 1A. **B.** Non-transgenic wild-type iL3s exposed to either histamine (+His) or a no-histamine control (-His) penetrated rat skin similarly. Bar graph shows the percentage of -His and +His wild-type iL3s that completed skin penetration. n = 15-16 iL3s per condition. ns = not significant, Fisher's exact test. **C.** Histamine exposure does not affect pushes and punctures. Violin plot depicts the percentage of time on skin that -His vs. +His wild-type iL3s spent engaging in pushes or punctures. n = 11-12 iL3s per condition. ns = not significant, Mann-Whitney test. The iL3s that had initiated penetration by the time the recording started were excluded from this analysis. **D.** Histamine treatment does not affect the time to first puncture. Violin plot shows the time taken by -His vs. +His wild-type iL3s to puncture the skin for the first time since placement on the skin surface. n = 15-16 iL3s per condition. ns = not significant, Mann-Whitney test. The dotted line at y = 5 indicates the time at which the assay ended. For C-D, dots depict individual worms, dashed lines indicate medians, and dotted lines indicate interquartile ranges. Behavioral parameters plotted in **B**-**D** were obtained from 3 independent replicate experiments.

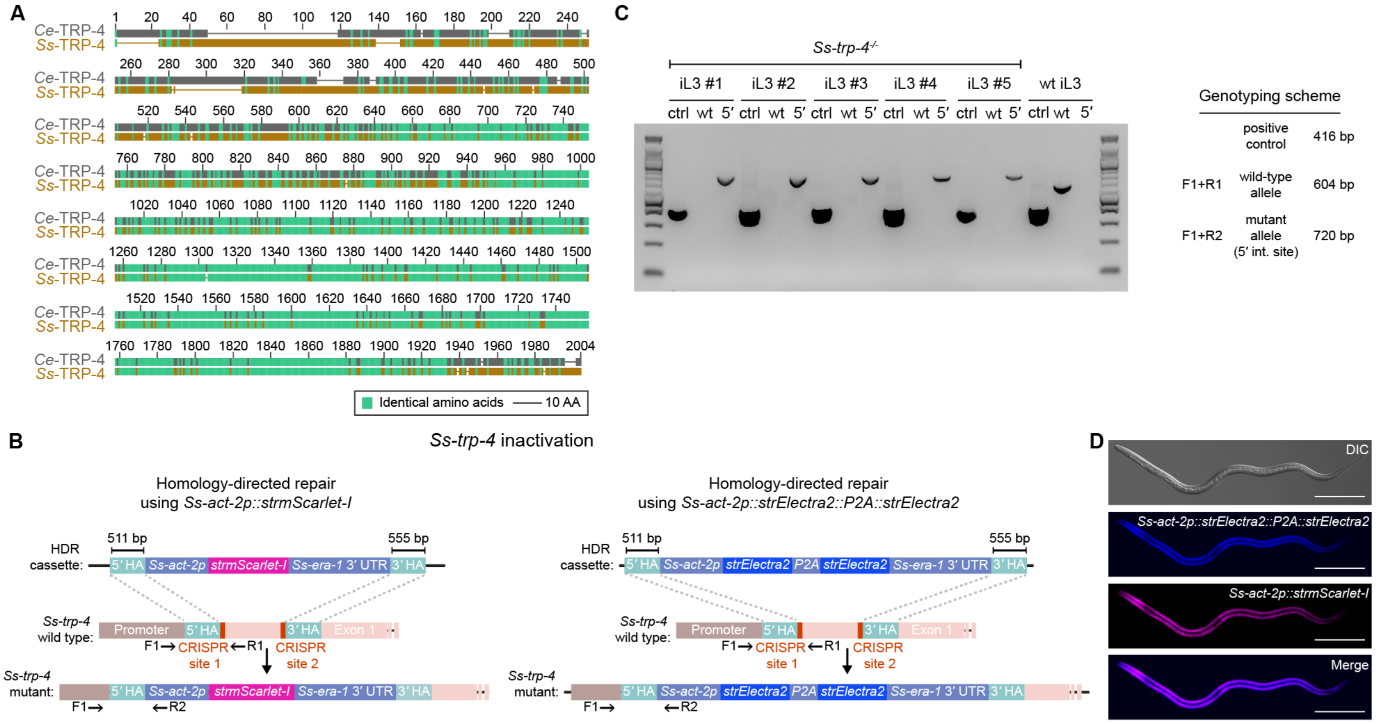


**Figure S7. Inactivation of *Ss*-TRP-4 using CRISPR/Cas9-mediated mutagenesis. A.** Schematic representation of the alignment of the amino acid sequences of the *C. elegans* TRP-4 protein (top) and the *S. stercoralis* TRP-4 protein (bottom). The amino acid sequences of the two proteins are 57.1% identical. Identical amino acids are depicted in green. Drawings are to scale and the scale bar = 10 amino acids (AA). **B.** Inactivation of *Ss-trp-4* was achieved by integration of a transcriptional reporter for the *Ss-act-2* gene at cut sites generated by Cas9. Two distinct sgRNAs, one targeting CRISPR site 1 and the other targeting CRISPR site 2, were used in conjunction with Cas9 to generate double-strand breaks at the *Ss-trp-4* locus; an *Ss-act-2* transcriptional reporter was then inserted between these two sites by homology-directed repair. The *act-2* transcriptional reporter consisted of the promoter of the *Ss-act-2* gene fused with a *Strongyloides*-codon-optimized gene encoding either mScarlet-I (left) or Electra2::P2A::Electra2[24] (right) and the *Ss-era-1* 3′ UTR. This transgene was flanked by 5′ and 3′ homology arms (HAs); the 5′ HA matched the 511 bp fragment immediately upstream of CRISPR site 1 in *Ss-trp-4* (depicted in red) and the 3′ HA matched a 555 bp fragment immediately downstream of CRISPR site 2 (red) in the same gene. Homology-directed repair following double-stranded breaks generated by Cas9 resulted in insertion of either the *Ss-act-2p::strmScarlet-I::Ss-era-1* 3′ UTR transgene or the *Ss-act-2p::strElectra2::P2A::strElectra2::Ss-era-1* 3′ UTR transgene into the *Ss-trp-4* locus, creating a stop codon early in the first exon and thereby preventing expression of the mutant allele. The approximate binding sites of the genotyping primers are also shown. The presence of a PCR amplicon from F1 and R1 indicated a wild-type locus, as R1 overlaps the region that is excised by Cas9. The presence of a PCR amplicon from F1 and R2 indicated a mutant locus, as F1 lies in the region of the *Ss-trp-4* promoter that is upstream of the 5′ HA and R2 lies in the *Ss-act-2* promoter. **C.** Image shows a representative agarose gel that was loaded and run with amplicons from genotyping PCRs of five *Ss-trp-4*$^{-/-}$ dual-colored iL3s and one wild-type iL3. For each worm, products from the following PCR reactions were loaded in order: a positive control for the PCR reaction (exon 1 of the *Ss-act-2* gene), which produced a 416 bp band (ctrl); a PCR reaction for genotyping the wild-type allele of *Ss-trp-4* (using primers F1 and R1), which produced a 604 bp band (wt) in wild-type iL3s; and a PCR reaction for genotyping the 5′ integration site of the HDR template (using primers F1 and R2), which produced a 720 bp band (5′) in carriers of the mutant *Ss-trp-4* allele. The first and last lanes carried a New England Biolabs 100 bp ladder. "5′ int. site" = 5′ integration site. **D.** Expression of the *Ss-act-2p::strElectra2::P2A::strElectra2::Ss-era-1* 3′ UTR (blue) and *Ss-act-2p::strmScarlet-I::Ss-era-1* 3′ UTR (magenta) transgenes over the entire body wall muscle of a dual-colored *Ss-trp-4* mutant. The worm is oriented with the ventral side facing downwards and the head to the left. Scale bar = 100 µm.

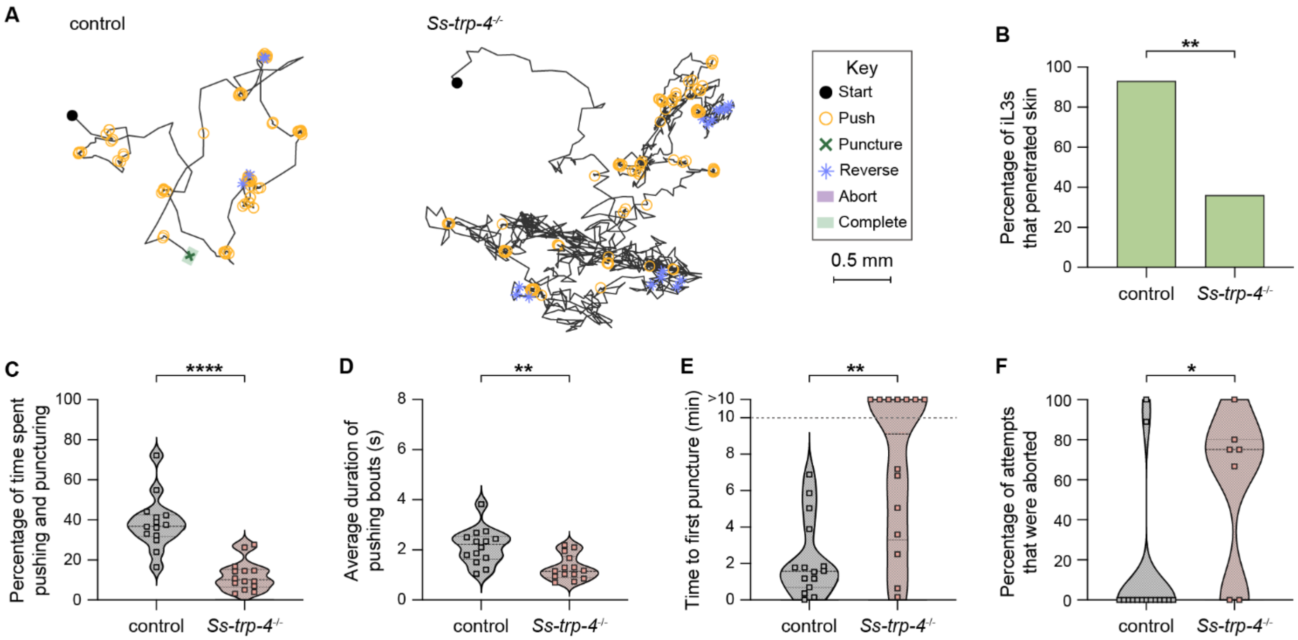


**Figure S8. Inactivation of *Ss-trp-4* severely impairs the ability to penetrate human skin. A.** *Ss-trp-4$^{-/-}$* iL3s exhibit reduced skin penetration and altered behaviors on human skin. Tracks show the behaviors of a representative wild-type iL3 that punctured and completed penetration and a representative *Ss-trp-4$^{-/-}$* iL3 that neither punctured nor completed penetration on human skin. Representative worms were defined as in Fig. 3A. The key details the behavioral motifs that were tracked. **B.** *Ss-trp-4$^{-/-}$* iL3s have severely reduced skin-penetration ability. Bar graphs show the percentage of wild-type and *Ss-trp-4$^{-/-}$* iL3s that completed skin penetration. n = 14-15 iL3s per genotype. $^{**}p<0.01$, Fisher's exact test. **C.** *Ss-trp-4$^{-/-}$* iL3s pushed and punctured the skin for less time than control worms. Violin plot depicts the percentage of time on skin that control and *Ss-trp-4$^{-/-}$* iL3s spent engaging in pushes or punctures. n = 14 iL3s per genotype. $^{****}p<0.0001$, unpaired t-test. The iL3s that had initiated penetration by the time the recording started were excluded from this analysis. **D.** The pushing bouts of *Ss-trp-4$^{-/-}$* iL3s were shorter than those of control iL3s. For each worm, the duration of each individual pushing bout was averaged and then plotted. n = 14 iL3s per genotype. $^{**}p<0.01$, unpaired t-test. **E.** Inactivation of *Ss-trp-4* inhibits punctures. Violin plot depicts the time taken by control vs. *Ss-trp-4$^{-/-}$* iL3s to puncture the skin for the first time since placement on skin. n = 14 iL3s per genotype. $^{**}p<0.01$, Mann-Whitney test. The dotted line at y = 10 indicates the time at which the assay ended; the dots above this line indicate animals that failed to puncture the skin by the end of the assay period. **F.** *Ss-trp-4$^{-/-}$* iL3s frequently abort penetration attempts. Violin plot depicts the percentage of penetration attempts, as defined by instances that the worm punctured and partially entered the skin, that were aborted. n = 7-15 iL3s per genotype. $^{*}p<0.05$, Mann-Whitney test. For **C-F**, dots depict individual worms, dashed lines indicate medians, and dotted lines indicate interquartile ranges. Behavioral parameters plotted in **B-F** were obtained from 3 independent replicate experiments, each using skin from a distinct human donor.

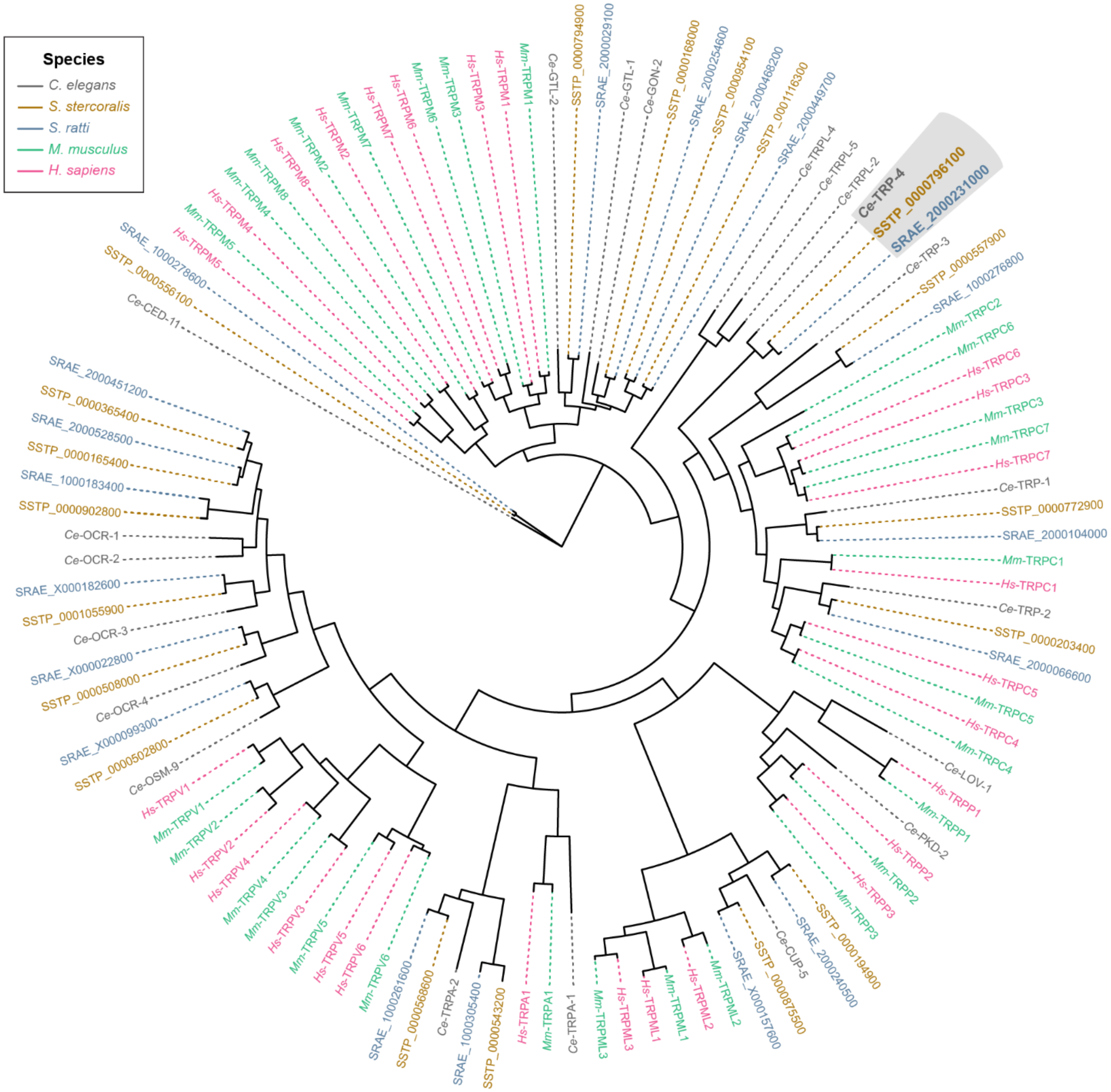


**Figure S9. *Ss*-TRP-4 is not conserved to humans.** Phylogenetic analysis of the TRP family members of *C. elegans* (gray), *S. stercoralis* (brown), *S. ratti* (blue), *Mus musculus* (green) and *Homo sapiens* (pink). The putative TRP family members of *S. stercoralis* and *S. ratti* were identified by performing TBLASTN searches of the *C. elegans* TRP channels[25-27] against either the *S. stercoralis* genome or the *S. ratti* genome in WBPS18. The sequences of known *M. musculus* and *H. sapiens* TRP channels[28] were retrieved from UniProtKB (Release 2024_06); the related accession numbers are listed in Supplemental Table 2.

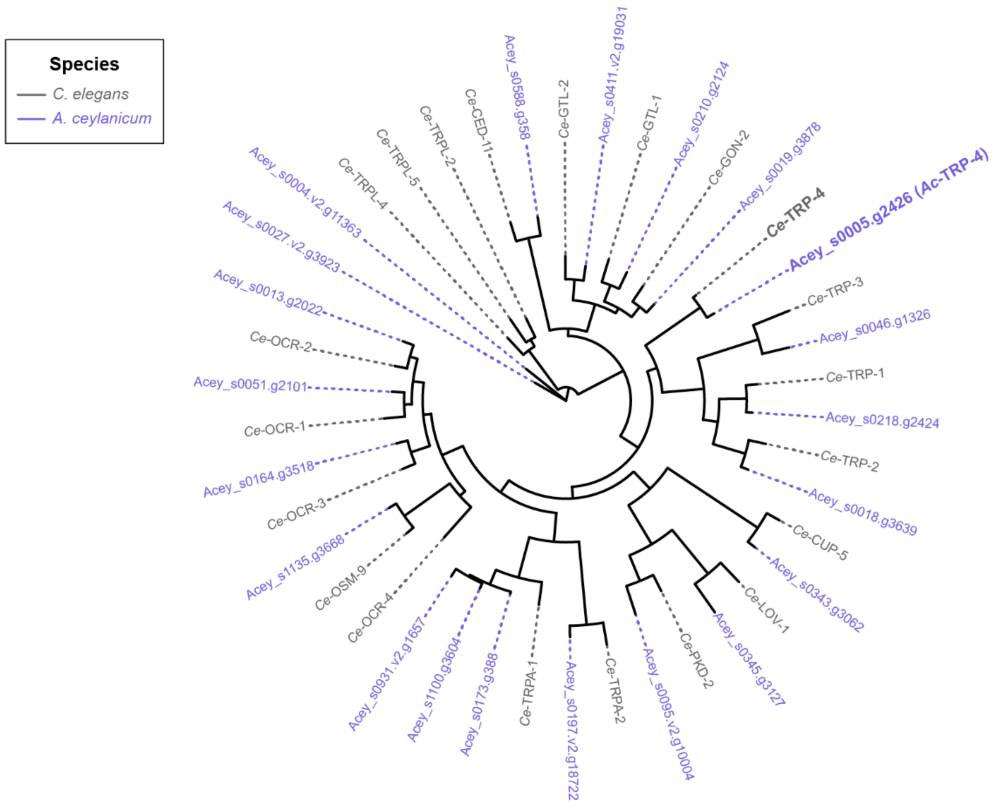


**Figure S10. *Ss*-TRP-4 is conserved to hookworms.** Phylogenetic analysis of the TRP family members of *C. elegans* (gray) and *A. ceylanicum* (purple). The putative TRP family members of *A. ceylanicum* were identified by performing TBLASTN searches of the *C. elegans* TRP channels[25-27] against the *A. ceylanicum* genome (PRJNA23179) in WBPS19.

### SUPPLEMENTAL TABLES

**Table S1.** Oligonucleotides used for genotyping worms from CRISPR/Cas9 experiments.

| Primer name | Sequence | Description/Use |
|---|---|---|
| SG78 | 5′-GTATTCCCTTCTATTGTTGGAAGACC-3′ | Forward primer for genotyping *Ss-act-2* exon 1; positive control |
| SG80 | 5′-CCTTCATAGATTGGTACAGTGTGAG-3′ | Reverse primer for genotyping *Ss-act-2* exon 1; positive control |
| RP32 | 5′-CGTAATTAAAAATATTTAAAAGTTACTATTACG-3′ | Forward primer in *Ss-cat-2* promoter for genotyping wild-type *Ss-cat-2* allele; “Primer F1” |
| RP33 | 5′-GGTATCTAATTTTGCTAATGGTGG-3′ | Reverse primer in *Ss-cat-2* exon 2 (overlapping CRISPR site) for genotyping wild-type *Ss-cat-2* allele; “Primer R1” |
| RP30 | 5′-CGGTATATTTTACTTCAATGTGG-3′ | Forward primer in *Ss-era-1* 3′ UTR for genotyping 3′ integration band; “Primer F2” |
| RP34 | 5′-TTAGCCATTTTATTTAACTCTTCACG-3′ | Reverse primer in *Ss-cat-2* exon 2 for genotyping *Ss-cat-2* 3′ integration band; “Primer R2” |
| RP39 | 5′-ATTTGGATGAAGACTTTTTCATACC-3′ | Forward primer in *Ss-trp-4* promoter for genotyping wild-type *Ss-trp-4* allele and 5′ integration band; “Primer F1” |
| RP46 | 5′-AATGATGACCATATTGCTCCAGC-3′ | Reverse primer in *Ss-trp-4* exon 1 for genotyping wild-type *Ss-trp-4* allele; “Primer R1” |
| RP28 | 5′-CGAGGAACCTCTTTTCCACAC-3′ | Reverse primer in *Ss-act-2* promoter for genotyping *Ss-trp-4* 5′ integration band: “Primer R2” |

**Table S2.** UniProtKB accession numbers for *M. musculus* and *H. sapiens* TRP channels.

| Species | Gene Name | UniProtKB accession number |
|---|---|---|
| *Homo sapiens* | TRPV1 | Q8NER1 |
| | TRPV2 | Q9Y5S1 |
| | TRPV3 | Q8NET8 |
| | TRPV4 | Q9HBA0 |
| | TRPV5 | Q9NQA5 |
| | TRPV6 | Q9H1D0 |
| | TRPA1 | O75762 |
| | TRPML1 | Q9GZU1 |
| | TRPML2 | Q8IZK6 |
| | TRPML3 | Q8TDD5 |
| | TRPP1 | P98161 |
| | TRPP2 | Q13563 |
| | TRPP3 | Q9P0L9 |
| | TRPC1 | P48995 |
| | TRPC3 | Q13507 |
| | TRPC4 | Q9UBN4 |
| | TRPC5 | Q9UL62 |
| | TRPC6 | Q9Y210 |
| | TRPC7 | Q9HCX4 |
| | TRPM1 | Q7Z4N2 |
| | TRPM2 | O94759 |
| | TRPM3 | Q9HCF6 |
| | TRPM4 | Q8TD43 |
| | TRPM5 | Q9NZQ8 |
| | TRPM6 | Q9BX84 |
| | TRPM7 | Q96QT4 |
| | TRPM8 | Q7Z2W7 |
| *Mus musculus* | TRPV1 | Q704Y3 |
| | TRPV2 | Q9WTR1 |
| | TRPV3 | Q8K424 |
| | TRPV4 | Q9EPK8 |
| | TRPV5 | P69744 |
| | TRPV6 | Q91WD2 |
| | TRPA1 | Q8BLA8 |
| | TRPML1 | Q99J21 |
| | TRPML2 | Q8K595 |
| | TRPML3 | Q8R4F0 |

| | | |
|---|---|---|
| | TRPP1 | O08852 |
| | TRPP2 | O35245 |
| | TRPP3 | A2A259 |
| | TRPC1 | Q61056 |
| | TRPC2 | Q9R244 |
| | TRPC3 | Q9QZC1 |
| | TRPC4 | Q9QUQ5 |
| | TRPC5 | Q9QX29 |
| | TRPC6 | Q61143 |
| | TRPC7 | Q9WVC5 |
| | TRPM1 | Q2TV84 |
| | TRPM2 | Q91YD4 |
| | TRPM3 | J9SQF3 |
| | TRPM4 | Q7TN37 |
| | TRPM5 | Q9JJH7 |
| | TRPM6 | Q8CIR4 |
| | TRPM7 | Q923J1 |
| | TRPM8 | Q8R4D5 |

### SUPPLEMENTAL MOVIE LEGENDS

**Movie S1. Skin-penetration behavior of an *S. stercoralis* iL3 on rat skin.** Movie shows time-lapse images of an *S. stercoralis* iL3 (expressing *Ss-act-2::strmScarlet-I*) penetrating rat skin. Images were acquired at 2 frames/s and the playback speed is 8 frames/s.

**Movie S2. Skin-penetration behavior of an *S. ratti* iL3 on rat skin.** Movie shows time-lapse images of an *S. ratti* iL3 (stained with DiI) penetrating rat skin. This iL3 was a vehicle-only control from the experiments with haloperidol. The point where the iL3 was initially placed on the skin is marked by an accumulation of the fluorescent dye. Images were acquired at 2 frames/s and the playback speed is 8 frames/s.

### SUPPLEMENTAL REFERENCES


1 Viney, M. E. & Lok, J. B. The biology of *Strongyloides* spp. *WormBook*, 1-17 (2015). https://doi.org:10.1895/wormbook.1.141.2

2 Castelletto, M. L. *et al.* Diverse host-seeking behaviors of skin-penetrating nematodes. *PLoS Pathog* **10**, e1004305 (2014). https://doi.org:10.1371/journal.ppat.1004305

3 Noskova, E. *et al.* *Strongyloides* in non-human primates: significance for public health control. *Philos Trans R Soc Lond B Biol Sci* **379**, 20230006 (2024). https://doi.org:10.1098/rstb.2023.0006

4 Genta, R. M. & Gomes, M. C. in Strongyloidiasis: a major roundworm infection of man (ed D. I. Grove) 105-132 (Taylor & Francis, 1989).

5 Dionisio, D. *et al.* *Strongyloides stercoralis*: ultrastructural study of newly hatched larvae within human duodenal mucosa. *J Clin Pathol* **53**, 110-116 (2000). https://doi.org:10.1136/jcp.53.2.110

6 Bryant, A. S. *et al.* A critical role for thermosensation in host seeking by skin-penetrating nematodes. *Curr Biol* **28**, 2338-2347 e2336 (2018). https://doi.org:10.1016/j.cub.2018.05.063

7 Gordon, C. A. *et al.* Strongyloidiasis. *Nat Rev Dis Primers* **10**, 6 (2024). https://doi.org:10.1038/s41572-023-00490-x

8 Hall, A. D., Luckett, K. M. & Williams, K. M. Bullous cutaneous larva migrans of the foot. *Am J Trop Med Hyg* **110**, 625-626 (2024). https://doi.org:10.4269/ajtmh.23-0750

9 Green, R., Somayaji, R. & Chia, J. C. Bullous cutaneous larva migrans. *CMAJ* **195**, E1040 (2023). https://doi.org:10.1503/cmaj.230583

10 Gomez-Moyano, E., Pilar, L. M., Simonsen, S. B. & Vera-Casano, A. A serpiginous, itchy rash on the foot. *Cleve Clin J Med* **83**, 494-495 (2016). https://doi.org:10.3949/ccjm.83a.15110

11 Lee, Y. & Hwang, K. Skin thickness of Korean adults. *Surg Radiol Anat* **24**, 183-189 (2002). https://doi.org:10.1007/s00276-002-0034-5

12 Oltulu, P., Ince, B., Kokbudak, N., Findik, S. & Kilinc, F. Measurement of epidermis, dermis, and total skin thicknesses from six different body regions with a new ethical histometric technique. *Turkish Journal of Plastic Surgery* **26**, 56-61 https://doi.org:10.4103/tjps.TJPS_2_17

13 Fitzpatrick, P. F. The aromatic amino acid hydroxylases: structures, catalysis, and regulation of phenylalanine hydroxylase, tyrosine hydroxylase, and tryptophan hydroxylase. *Arch Biochem Biophys* **735**, 109518 (2023). https://doi.org:10.1016/j.abb.2023.109518

14 Banerjee, N. *et al.* Carbon dioxide shapes parasite-host interactions in a human-infective nematode. *Curr Biol* (2024).

15 Gang, S. S. *et al.* Targeted mutagenesis in a human-parasitic nematode. *PLoS Pathog* **13**, e1006675 (2017). https://doi.org:10.1371/journal.ppat.1006675

16 Ashton, F. T., Zhu, X., Boston, R., Lok, J. B. & Schad, G. A. *Strongyloides stercoralis*: amphidial neuron pair ASJ triggers significant resumption of development by infective larvae under host-mimicking *in vitro* conditions. *Exp Parasitol* **115**, 92-97 (2007). https://doi.org:10.1016/j.exppara.2006.08.010

17 Stoltzfus, J. D., Massey, H. C., Jr., Nolan, T. J., Griffith, S. D. & Lok, J. B. *Strongyloides stercoralis* age-1: a potential regulator of infective larval development in a parasitic nematode. *PLoS One* **7**, e38587 (2012). https://doi.org:10.1371/journal.pone.0038587

18 Stoltzfus, J. D., Bart, S. M. & Lok, J. B. cGMP and NHR signaling co-regulate expression of insulin-like peptides and developmental activation of infective larvae in *Strongyloides stercoralis*. *PLoS Pathog* **10**, e1004235 (2014). https://doi.org:10.1371/journal.ppat.1004235

19 Gang, S. S. *et al.* Chemosensory mechanisms of host seeking and infectivity in skin-penetrating nematodes. *Proc Natl Acad Sci USA* **117**, 17913-17923 (2020). https://doi.org:10.1073/pnas.1909710117

20 Patel, R. *et al.* The generation of stable transgenic lines in the human-infective nematode *Strongyloides stercoralis*. *G3 (Bethesda)* **14** (2024). https://doi.org:10.1093/g3journal/jkae122

21 Pokala, N., Liu, Q., Gordus, A. & Bargmann, C. I. Inducible and titratable silencing of *Caenorhabditis elegans* neurons in vivo with histamine-gated chloride channels. *Proc Natl Acad Sci USA* **111**, 2770-2775 (2014). https://doi.org:10.1073/pnas.1400615111

22 Bryant, A. S., Ruiz, F., Lee, J. H. & Hallem, E. A. The neural basis of heat seeking in a human-infective parasitic worm. *Curr Biol* **32**, 2206-2221 e2206 (2022). https://doi.org:10.1016/j.cub.2022.04.010

23 Castelletto, M. L. & Hallem, E. A. Generating transgenics and knockouts in *Strongyloides* species by microinjection. *J Vis Exp* (2021). https://doi.org:10.3791/63023

24 Papadaki, S. *et al.* Dual-expression system for blue fluorescent protein optimization. *Sci Rep* **12**, 10190 (2022). https://doi.org:10.1038/s41598-022-13214-0
25 Xiao, R. & Xu, X. Z. *C. elegans* TRP channels. *Adv Exp Med Biol* **704**, 323-339 (2011). https://doi.org:10.1007/978-94-007-0265-3_18
26 Kahn-Kirby, A. H. & Bargmann, C. I. TRP channels in *C. elegans*. *Annu Rev Physiol* **68**, 719-736 (2006). https://doi.org:10.1146/annurev.physiol.68.040204.100715
27 Goodman, M. B. Mechanosensation. *WormBook*, 1-14 (2006). https://doi.org:10.1895/wormbook.1.62.1
28 Venkatachalam, K. & Montell, C. TRP channels. *Annu Rev Biochem* **76**, 387-417 (2007). https://doi.org:10.1146/annurev.biochem.75.103004.142819
